# Heart rate dynamics embed a shared representation of individual brain organization and cognitive function

**DOI:** 10.64898/2026.08.04.742623

**Authors:** Liyi Qian, Ben D. Fulcher, Sihan Ju, Zhongyi Xiao, Yuhao Xu, Yunman Xia, Yanqun Zheng, Liang Chen, Yunzeng Zou, Gunter Schumann, Xiang Zou, Kun Song, Ying Mao

## Abstract

Idiosyncratic brain functional organization shapes intricate cardiac dynamics through central-peripheral autonomic interactions, yet a comprehensive mapping between multi-scale heart rate dynamics and whole-brain functional architecture remains lacking. Here, combining highly comparative time-series analysis with multi-modal neuroimaging and intracranial electrophysiology, we establish heart rate dynamics as a physiological fingerprint that maps onto whole-brain functional architecture. Partial least squares analysis revealed a generalizable latent axis linking reduced heart rate temporal complexity and elevated micro-scale predictability to heightened resting-state functional connectivity across default mode, salience, and sensorimotor networks. This covariance aligns spatially with serotonergic, noradrenergic, and cholinergic neuromodulatory gradients, persists across physiological confound controls and cross-session validations, derives support from human intracranial electrophysiological recordings, and extends to active cognitive states. Furthermore, brain-covarying heart rate signatures underpin the predictive capacity of heart rate dynamics for individual fluid and crystallized intelligence, demonstrating a shared representational substrate. Our findings demonstrate a robust neurovisceral coupling architecture, establishing well-characterized heart rate dynamics as a scalable, neurobiologically anchored window into human brain functional organization and cognitive traits.

## Introduction

Heart rate exhibits complex temporal fluctuations governed by the autonomic nervous system (ANS)^1,2^. The parasympathetic branch decreases heart rate and contributes predominantly to rapid, short-term variability, whereas the sympathetic branch increases heart rate and shapes slower temporal fluctuations^3,4^. Their dynamic interaction generates multi-scale cardiac activity^5,6^. In response to internal and external demands, the ANS interacts with the central autonomic network (CAN)^7,8^ to coordinate adaptive physiological responses, forming the basis of the neurovisceral integration model^9–11^. The CAN encompasses a hierarchical set of brain regions extending from the brainstem and subcortical structures to high-order cortical areas^12–15^. This central-peripheral organization suggests that stable individual differences in brain function and behavior may be reflected in heart rate dynamics. Because electrocardiography (ECG) and photoplethysmography (PPG) are widely available in clinical and ambulatory settings, comprehensive analysis of heart rate dynamics may provide an accessible physiological window into individual brain organization^16,17^.

Previous studies have identified associations of conventional time- and frequency-domain heart rate variability (HRV) measures^18,19^ with isolated brain features and behavioral phenotypes^20–23^. However, these measures capture only a restricted subset of the temporal structure contained in cardiovascular signals. Moreover, the brain operates as an integrated dynamical system, and a limited set of conventional HRV indices may not fully capture the landscape of brain-related cardiovascular dynamics. This limitation may arise from the analytical measures used rather than from the physiological signal itself, because heart rate signals contain a broader range of nonlinear, complexity, autocorrelative, and predictive properties^24,25^. Emerging studies suggest that conventional measures may incompletely capture complex neural dynamics^26–28^, whereas broader dynamical descriptors may provide complementary information in selected clinical and cognitive settings^29–31^. Nevertheless, whether heart rate dynamical properties are reproducibly associated with whole-brain functional organization remains uncertain. In addition, while large-scale functional neuroimaging cohorts now enable such high-dimensional feature space explorations, associations based on blood oxygen level-dependent (BOLD) functional magnetic resonance imaging may be influenced by respiratory and global vascular fluctuations^32–36^. Thus, it is critical to verify whether the linkage represents a genuine shared trait between the central and peripheral nervous systems, and whether it carries meaningful behavioral significance.

In this study, we sought to establish an integrated framework to map heart-brain-behavior associations. We applied highly comparative time-series analysis^37,38^ to extract thousands of quantitatively defined heart rate dynamical properties in a large cohort of healthy young adults. Using this expansive feature set, we first evaluated whether heart rate dynamical properties function as an individual physiological fingerprint. We then used partial least squares analysis to identify latent covariance between heart rate dynamics and resting-state functional connectivity. Robustness was assessed by independent cohort replication, adjustment for physiological confounders, cross-session projection, and validation using concurrent intracranial stereo-electroencephalography (sEEG) and electrocardiography. Expanding beyond spontaneous fluctuations, we further investigated the persistence and state-dependent shifts of this coupling during active task engagement. Finally, we examined whether the brain-covarying heart rate signatures predicted individual behavioral traits, as well as quantifying their representational alignment.

## Results

### Individual fingerprinting of heart rate dynamics

We extracted PPG signals recorded simultaneously with resting-state fMRI from the Human Connectome Project Young Adult (HCP-YA) dataset^39^ and estimated instantaneous heart rate (Fig. 1a). To comprehensively characterize individual heart rate dynamics, we constructed a unified high-dimensional profile by combining classic time-domain and frequency-domain HRV metrics with the highly comparative time-series analysis (hctsa) framework^37,38^, to transform temporally structured heart rate into a common feature space. Importantly, we refrain from referring to these combined descriptors merely as HRV. Instead, this comprehensive feature library encompasses not only conventional HRV measures and previously explored complexity and fractal metrics^5,40,41^ but also a broader array of dynamical properties, including not limited to forecasting measures, linear and non-linear autocorrelation properties, and models fitting performance. Following the removal of features with minimal variance or lacking meaningful interpretability, 6,817 features were retained, hereafter referred to as heart rate dynamical properties.

**Figure 1.**
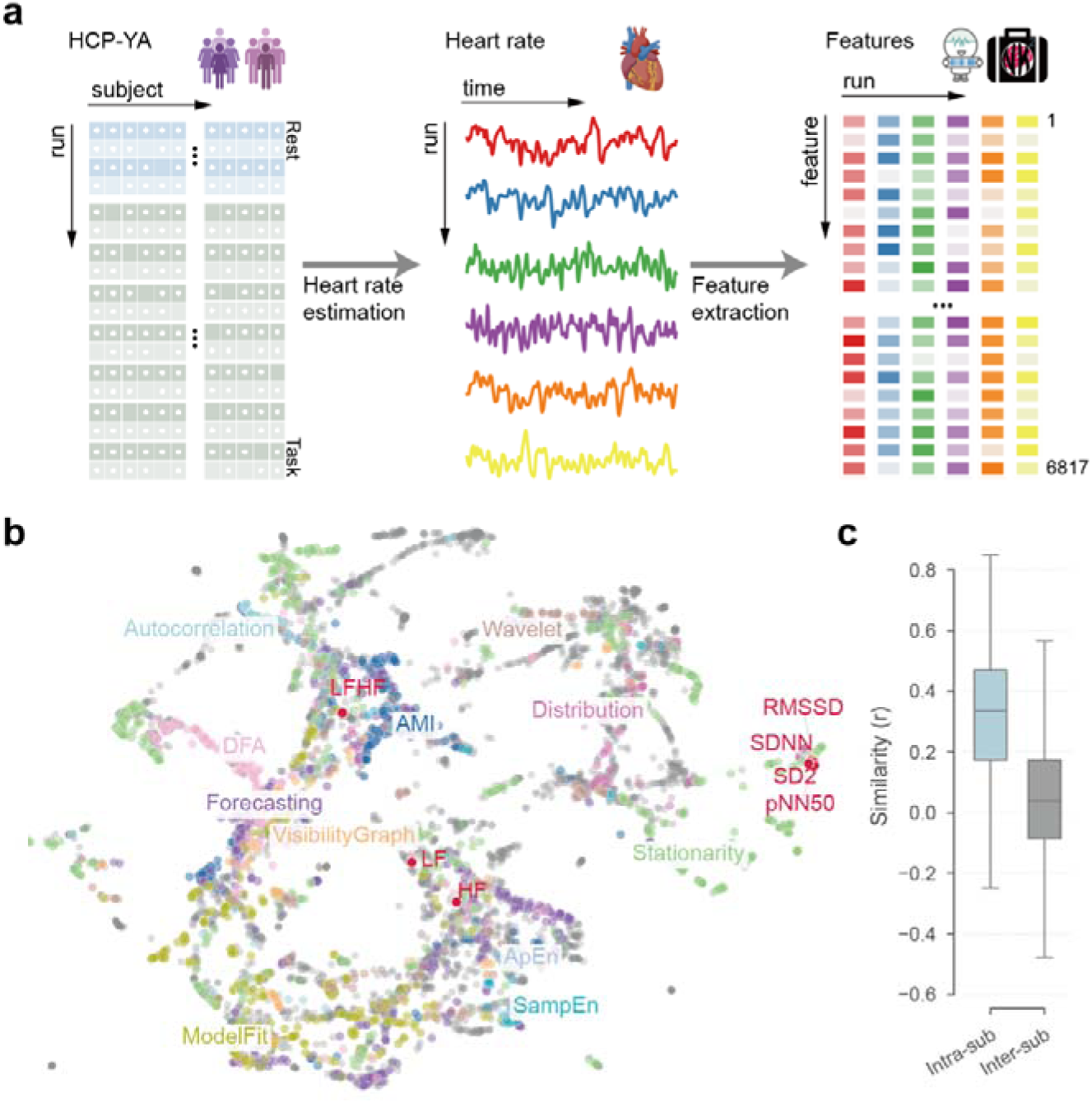
Characterization of individual heart rate dynamics. **a**, Feature extraction pipeline. Instantaneous heart rate time series are estimated from PPG recordings in the HCP-YA dataset. A unified high-dimensional profile comprising 6,817 features is extracted per run, combining classic HRV measures with the highly comparative time-series analysis (hctsa) framework. **b**, UMAP embedding of the joint feature space. Classic HRV metrics are annotated in red. The remaining hctsa-derived features are color-coded according to their broad algorithmic classes. **c**, Similarity distributions. Boxplots display the intra-subject and inter-subject similarities, evaluated using the Pearson correlation coefficient (*r*) of the feature vectors after robust-sigmoid normalization for each feature across runs.

To evaluate whether this strategy provides a more comprehensive perspective on heart rate dynamics, we mapped the joint feature space into a low-dimensional manifold using Uniform Manifold Approximation and Projection (UMAP) (Fig. 1b). The embedding demonstrates that traditional time-domain and frequency-domain HRV metrics occupy a constrained topological subspace, indicating a restricted capacity to resolve diverse temporal properties. The complexity and fractal descriptors, such as approximate entropy (ApEn), sample entropy (SampEn), and detrended fluctuation analysis (DFA), occupy distinct regions within the manifold, separating themselves from traditional HRV metrics. Furthermore, broader feature classes reflecting properties such as stationarity, forecasting, and autocorrelation distribute across the remaining representational space, capturing heart rate properties unaccounted for by standard HRV analysis.

Building upon this comprehensive characterization, we next evaluated whether these heart rate dynamical properties possess sufficient individual specificity to serve as a physiological fingerprint. To this end, we quantified individual discriminability of heart rate dynamical properties using the identifiability index^42,43^, which quantifies the difference between the average within-subject similarity and the average between-subjects similarity (Fig. 1c). An identifiability index ≥ 0.8 is typically considered large^43,44^. Among participants with four resting-state PPG recordings (*n* = 510), the identifiability index reached 1.53, indicating a strong capacity for the dynamical characteristics of heart rate to act as fingerprints for individual differentiation.

### Heart rate dynamics covary with intrinsic brain functional connectivity

Based on the individual specificity observed in heart rate dynamics, we sought to test whether these peripheral profiles could characterize individual-specific neural patterns. Since brain resting-state functional connectivity (RSFC) is a well-established neural fingerprint^43,45,46^, we first investigated whether heart rate dynamical properties map onto individual variations in RSFC.

Participants with at least one resting state session that passed quality control were included in the analysis (*n* = 878; see Methods for quality control details). We applied partial least squares (PLS) analysis to identify latent variables capturing maximal covariance between RSFC and heart rate dynamical properties (Fig. 2a). To mitigate potential overfitting due to excessive feature dimensionality, we employed a relatively coarse parcellation, generating a 116-node functional connectivity matrix comprising 100 cortical and 16 subcortical regions^47,48^. The resulting input matrices consisted of 6,817 heart rate features and 6,670 unique RSFC edges per participant. To control for potential confounds, we regressed out age, sex, and head motion from both matrices before PLS analysis.

**Figure 2.**
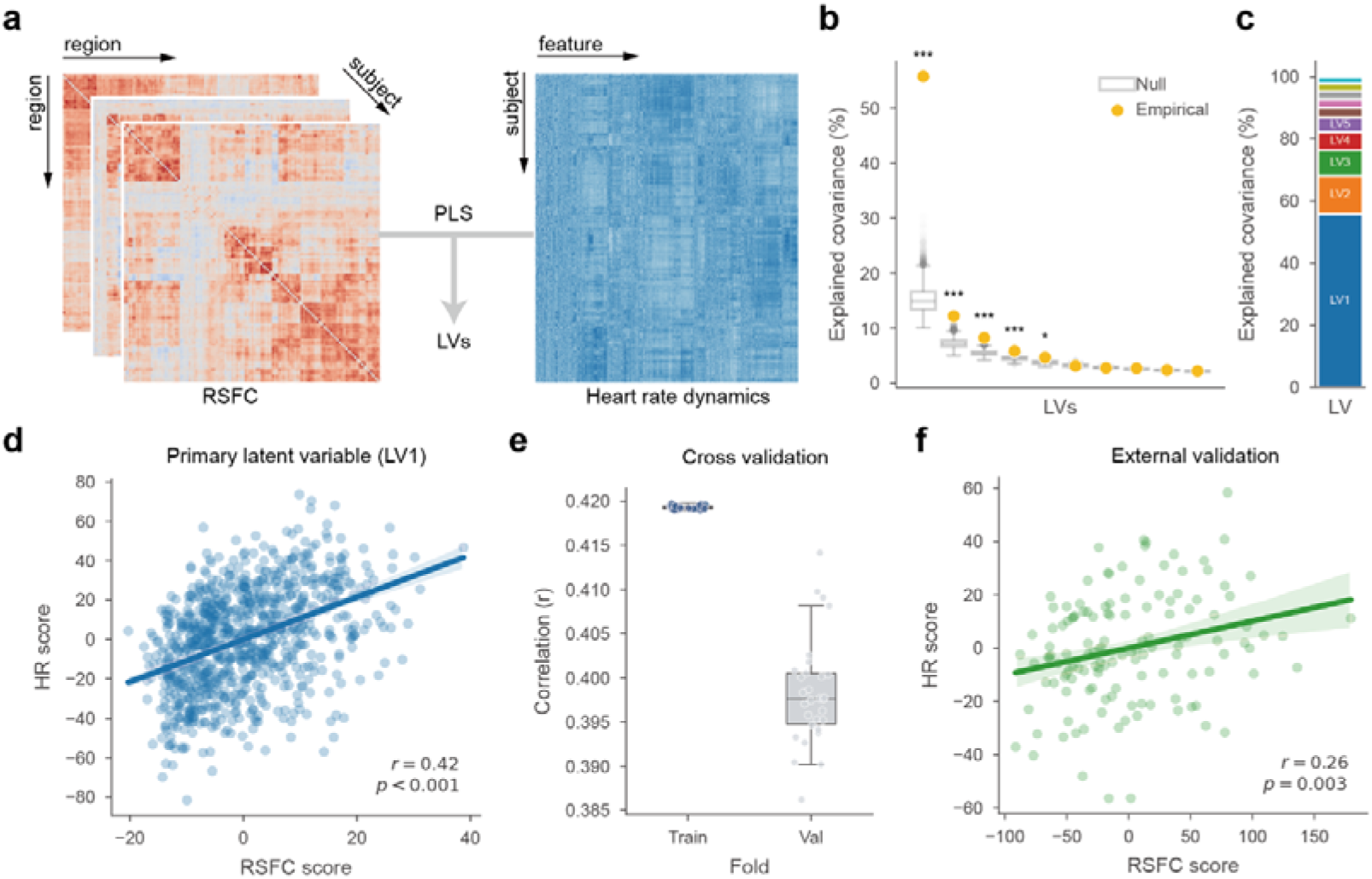
Linking heart rate dynamical properties to resting-state functional connectivity. **a**, Workflow of the PLS analysis mapping resting-state functional connectivity (RSFC) matrices to heart rate dynamical properties across subjects to identify latent variables (LVs). **b**, Explained covariance of LVs from empirical data (yellow dot) compared with a null distribution (gray box) obtained from 10,000 unrotated permutations (* *p* < 0.05, \*\**p* < 0.01, *** *p* < 0.001). **c**, Relative proportion of explained covariance for each identified LV. **d**, The correlation between individual RSFC scores and heart rate scores for LV1. The shaded area indicates a 95% confidence interval. *r* and *p* represent the Pearson correlation coefficient and the corresponding p-value, respectively. **e**, The distribution of correlation coefficients between RSFC and heart rate scores within the training and validation sets across 30 repeated 10-fold cross-validation. **f**, RSFC scores against heart rate scores in the external validation cohort. The shaded area indicates a 95% confidence interval. *r*, Pearson correlation coefficient.

PLS revealed a dominant latent variable (LV1) that accounted for 56% of the covariance between RSFC and heart rate dynamical properties (Fig. 2b, c). By projecting individual data onto the weight vectors corresponding to the LV1, we derived subject-specific RSFC and HR scores, which were positively correlated (*r* = 0.42, *p* < 0.001) (Fig. 2d).

In line with methodological recommendations^49^, we conducted 10,000 non-rotated permutations to impose a more stringent criterion on LV1, confirming its statistical significance (*p* < 0.0001). To assess the generalizability of this latent variable, we conducted 10-fold cross-validation. This latent variable remained robust across folds, yielding a mean training correlation of *r* = 0.42 and a mean validation correlation of *r* = 0.39 (Fig. 2e). Similar results were obtained using 216-, 376-, and 416- region parcellations, indicating that the identified covariance pattern was robust to spatial resolution (Supplementary Fig. 1).

To evaluate the generalizability across datasets, we introduced an independent cohort, LEMON dataset^50^ which differed in demographic characteristics and data acquisition protocols. A total of 132 participants were retained, comprising 97 males (73.5%) and 35 females (26.5%) with a median age of 25.0 years (IQR, 20.0–56.3). After data preprocessing, construction of RSFC and heart rate dynamical properties matrices, and regressing out confounds, we projected this data onto the HCP-YA derived LV1 weight vectors. This yielded a significant positive correlation between RSFC and HR scores (*r* = 0.26, *p* = 0.003) (Fig. 2f). In contrast, although the second latent variable also passed the permutation test in the HCP-YA dataset, it failed to replicate in this external cohort (*r* = 0.15, *p* = 0.08) and was therefore excluded from subsequent analyses.

For both modalities, feature contributions were indexed by loadings, defined as the Pearson correlation between each feature and corresponding PLS score^51^. Among heart rate dynamical properties, the most negative loadings were dominated by complexity measures, primarily spanning ApEn and SampEn (Fig. 3a; Supplementary Table 1). This pattern was accompanied by lower forecasting errors, such as predicting the next value from its immediate historical steps. The highest positive loadings were concentrated among properties derived from visibility graph analysis^52,53^, with the strongest loading reaching 0.93. Notably, these dominant features evaluate sub-second, micro-scale temporal structures (Supplementary Table 1). In comparison, conventional HRV measures which summarize macro-scale variability showed weaker loadings, including LF/HF ratio (0.47), SD2 (0.54), HF (-0.17), SDNN (0.44), LF (0.29), pNN50 (0.02), and RMSSD (0.12) (Supplementary Fig. 2). Taken together, higher heart rate scores were characterized by lower temporal complexity and higher predictability (Fig. 3b).

**Figure 3.**
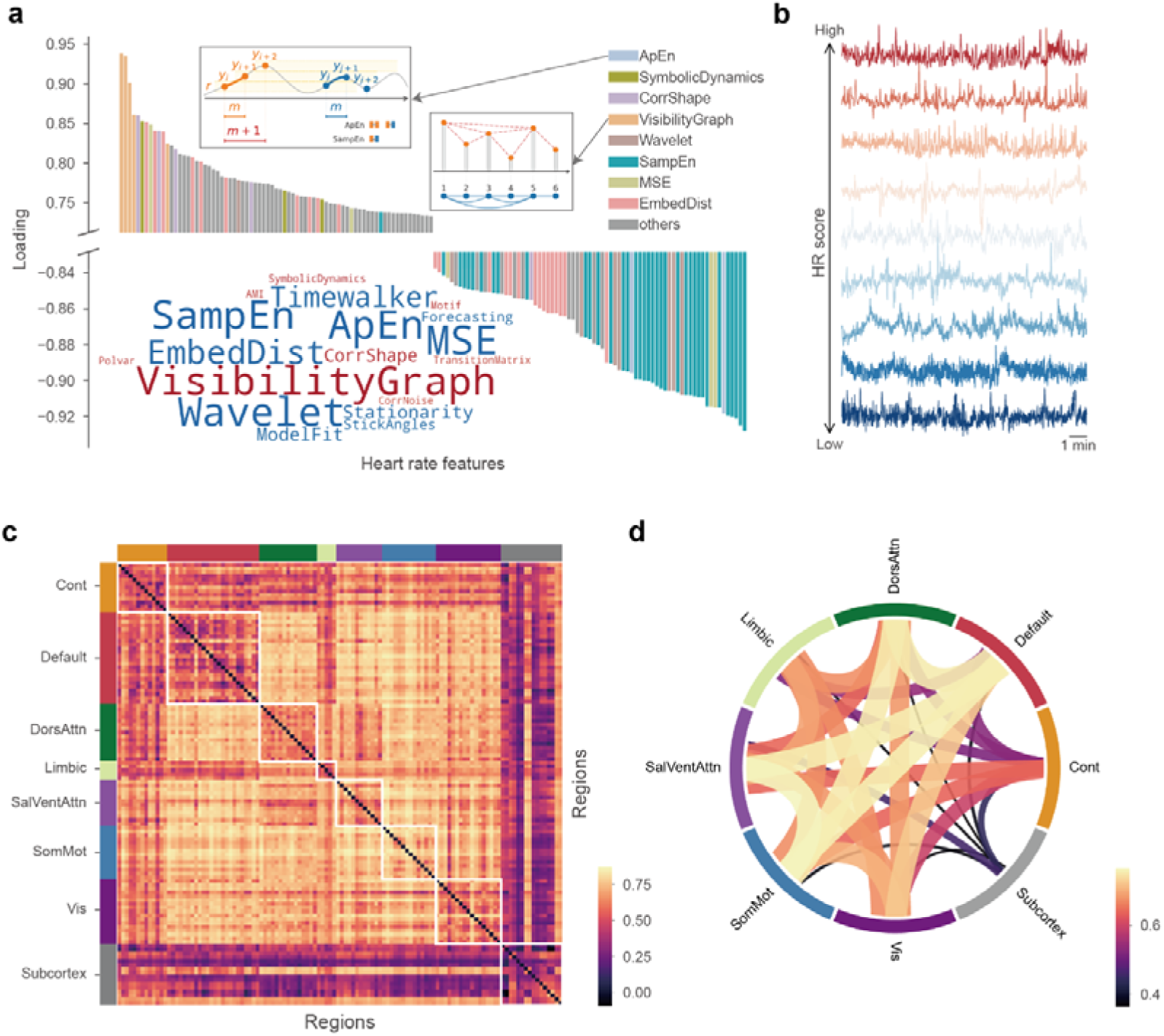
Covariation pattern between heart rate dynamics and resting-state functional connectivity. **a**, Loadings of heart rate features for the primary latent variable. Features are colored by their domains, with the inset word cloud illustrating the highest-loading features sized by their absolute loading values. **b**, Representative instantaneous heart rate time series ordered by their corresponding heart rate scores. **c**, The regional functional connectivity loadings across the brain connectome. Regions are organized and color-coded by Yeo-7 functional networks. **d**, The network-level RSFC loadings. The color of the connecting bands represents the magnitude of the network-level loading.

Turning to RSFC, we observed widespread positive loadings across the connectome (Fig. 3c). Specifically, the LV1 captured higher within-network connectivity of the somatomotor network (SomMot), as well as higher inter-network connectivity among the default mode (Default), dorsal attention (DorsAttn), salience/ventral attention (SalVentAttn), and somatomotor networks (Fig. 3c, d). By summing the connection loadings across all edges for each brain region, we identified the sensorimotor, insula, and prefrontal areas as the primary regional contributors to LV1 (Fig. 4a, b). To examine the potential neurobiological context of this cortical topography, we compared the regional loading map with 20 positron emission tomography (PET)-derived cortical distributions of neuromodulatory systems^54,55^. Spatial similarity was quantified using Spearman correlation with significance evaluated using 10,000 spatial autocorrelation-preserving permutations^56^ followed by FDR correction. Five neuromodulatory maps were significantly associated with the regional loading pattern (Fig. 4c). The strongest correlations were observed for the serotonin transporter (5HTT), vesicular acetylcholine transporter (VAChT), and norepinephrine transporter (NAT) (Fig. 4d-f). Additional associations were found for dopamine- and glutamate-related maps.

**Figure 4.**
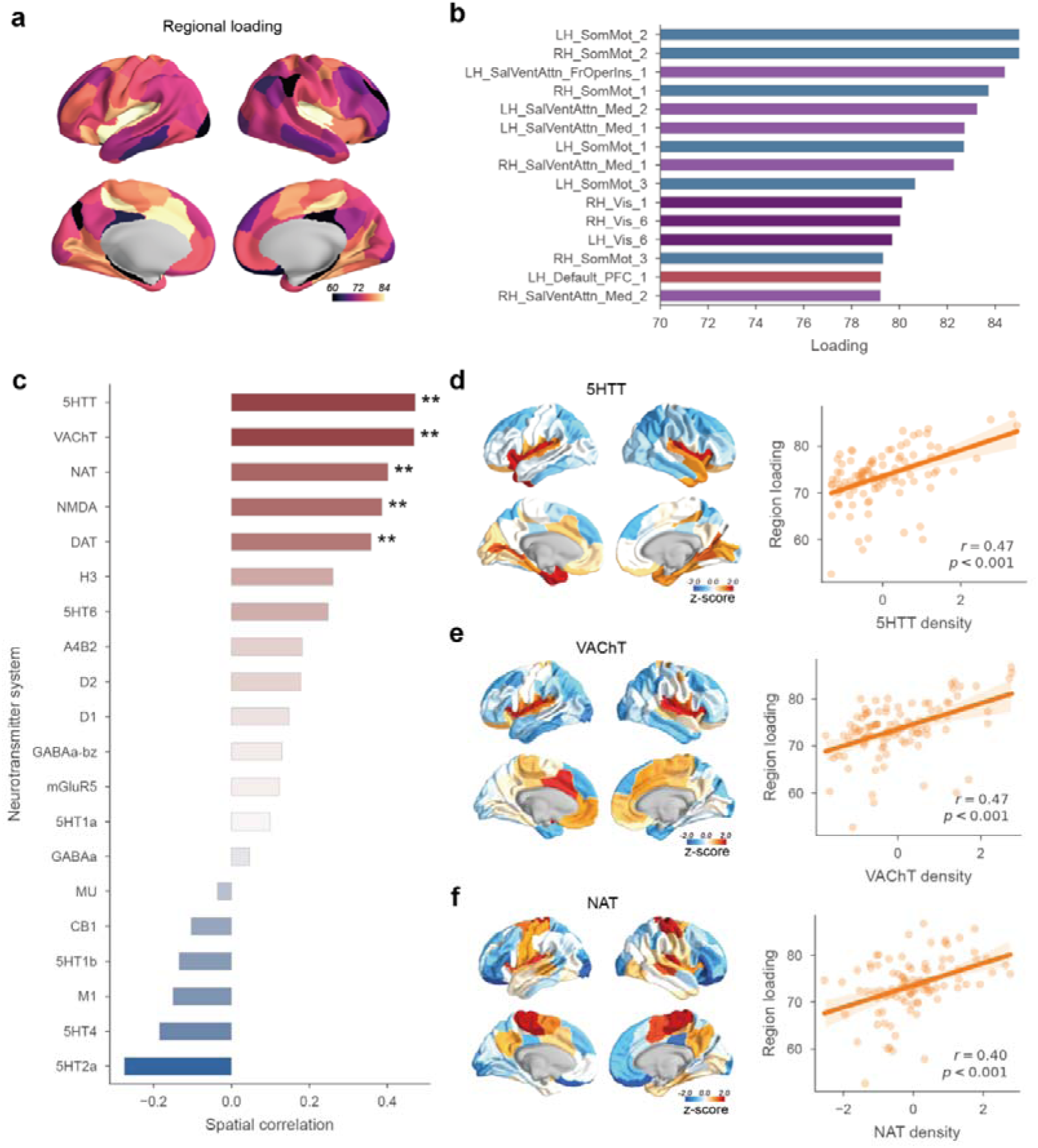
Cortical topography of regional loadings and spatial associations with neuromodulatory systems. **a**, Cortical surface mapping of the regional loadings, defined as the sum of its functional connectivity loadings. **b**, The top 15 cortical regions with the highest regional loading values, color-coded by their respective Yeo-7 functional networks. **c**, Spearman correlation between the regional loading maps and PET-derived cortical distributions of 20 neurotransmitter receptors and transporters. Significance was assessed using a spin test with 10,000 permutations (\*\**p* < 0.01). **d–f**, The three strongest neuromodulatory correlates: serotonin transporter (5-HTT) (d), vesicular acetylcholine transporter (VAChT) (e), and norepinephrine transporter (NAT) (f). Each panel displays the PET-based cortical density map (left) alongside regional loadings plotted against density across brain regions (right; shaded area indicates the 95% confidence interval).

Collectively, these findings delineate a robust and generalizable axis of heart-brain covariance, demonstrating that individual-specific heart rate dynamics map onto large-scale intrinsic neural network patterns anchored by specific neuromodulatory systems.

### Heart-brain covariance represents a stable neurophysiological trait

The central premise of linking heart rate dynamics to RSFC is to establish peripheral autonomic profiles as a biologically grounded, non-invasive window into individual variations in neural architecture. Although spatial alignment with neuromodulatory maps supported the neurobiological relevance of the identified heart-brain covariance, the BOLD signal remains an indirect proxy of neural activity and may be influenced by systemic physiological factors^57^. It is therefore critical to discern whether the individual RSFC variance captured by heart rate dynamical properties reflects a stable neurophysiological trait, or merely represents spurious hemodynamic alignments driven by systemic confounding factors.

We first addressed the potential confounding effects of respiration, which exhibits individual-specific characteristics and is known to modulate both heart rate^36^ and the BOLD signal^35^. After regressing out respiratory measures from the heart rate and the RSFC scores, the residual correlation remained significant and similar in magnitude to the primary result (*r* = 0.40, *p* < 0.001) (Supplementary Fig. 3), demonstrating that the observed covariance was not explained by these respiratory measures.

We next addressed the potential confounding by physiological co-fluctuations driven by internal or external cues. Recent evidence indicates that the fMRI global signal tightly co-fluctuates with peripheral physiological rhythms^33^, which could mimic individual-level covariance even in the absence of trait-like specificity. To explicitly test whether our identified covariance is driven solely by this global physiological state, we re-evaluated our PLS analysis after global signal regression (GSR). We assessed this robustness through two complementary steps. First, we directly projected the global signal regression (GSR) preprocessed RSFC matrices onto the LV1 weight vectors derived from the non-GSR data. Although the correlation between RSFC and HR scores was expectedly attenuated after removing global variance, it remained statistically significant (*r* = 0.27, *p* < 0.001), suggesting that the core spatial topography of the covariation is largely preserved (Supplementary Fig. 4a). Second, an independent PLS analysis using the GSR-preprocessed features again identified a significant latent variable (Supplementary Fig. 4b). Thus, although global co-fluctuation partially contributes to the observed covariance, it did not fully account for the identified heart-brain association.

We further reasoned that if this covariance reflected a stable individual trait rather than instantaneous temporal synchronization, the coupling should remain detectable when neural and cardiac data are acquired non-concurrently. To evaluate cross-session stability, we analyzed participants with multiple resting-state sessions (*n* = 837) and paired their RSFC matrices with heart rate dynamical properties extracted from non-concurrent runs (Supplementary Fig. 4c). Projection of these temporally segregated data pairs onto the original LV1 weight vectors derived from the primary synchronous analysis yielded significant correlations in both cross-paired configurations (*r* = 0.29 and *r* = 0.33, both *p* < 0.001) (Supplementary Fig. 4d). These findings support the cross-session stability of the observed heart–brain covariance and argue against an explanation based solely on transient temporal synchronization.

To determine whether the identified covariance was supported by direct neural electrophysiological recordings, we further evaluated it using concurrent stereo-electroencephalography (sEEG) and electrocardiography (ECG) (Fig. 5a). A total of 41 patients were included, comprising 23 males (56.1%) and 18 females (43.9%), with a median age of 24.0 years (IQR, 17.0–31.0). Each patient had 54 to 328 recording sites, which were anatomically mapped to 116 brain regions. For each patient, we extracted simultaneous resting-state sEEG and ECG segments that were free of epileptiform activity. We then computed the gamma-band envelope for each site, which has been proposed to reflect a neural component underlying the BOLD signal^58–60^. Functional connectivity was estimated using Pearson correlations between regional high-gamma envelopes. Due to the limited spatial coverage of sEEG within individual patients, the analysis was restricted to region pairs sampled in at least 10 patients. We next projected each patient’s heart rate dynamical properties onto the HCP-derived LV1 heart rate weight vector to obtain an individual heart rate score. Finally, we evaluated the relationship between FC edges and these individual HR scores by calculating their Pearson correlations. As shown in Fig. 5b, sEEG-derived functional connectivity predominantly displayed positive correlations with individual HR scores, corroborating the phenomena identified in our fMRI data analysis at an electrophysiological level.

**Figure 5.**
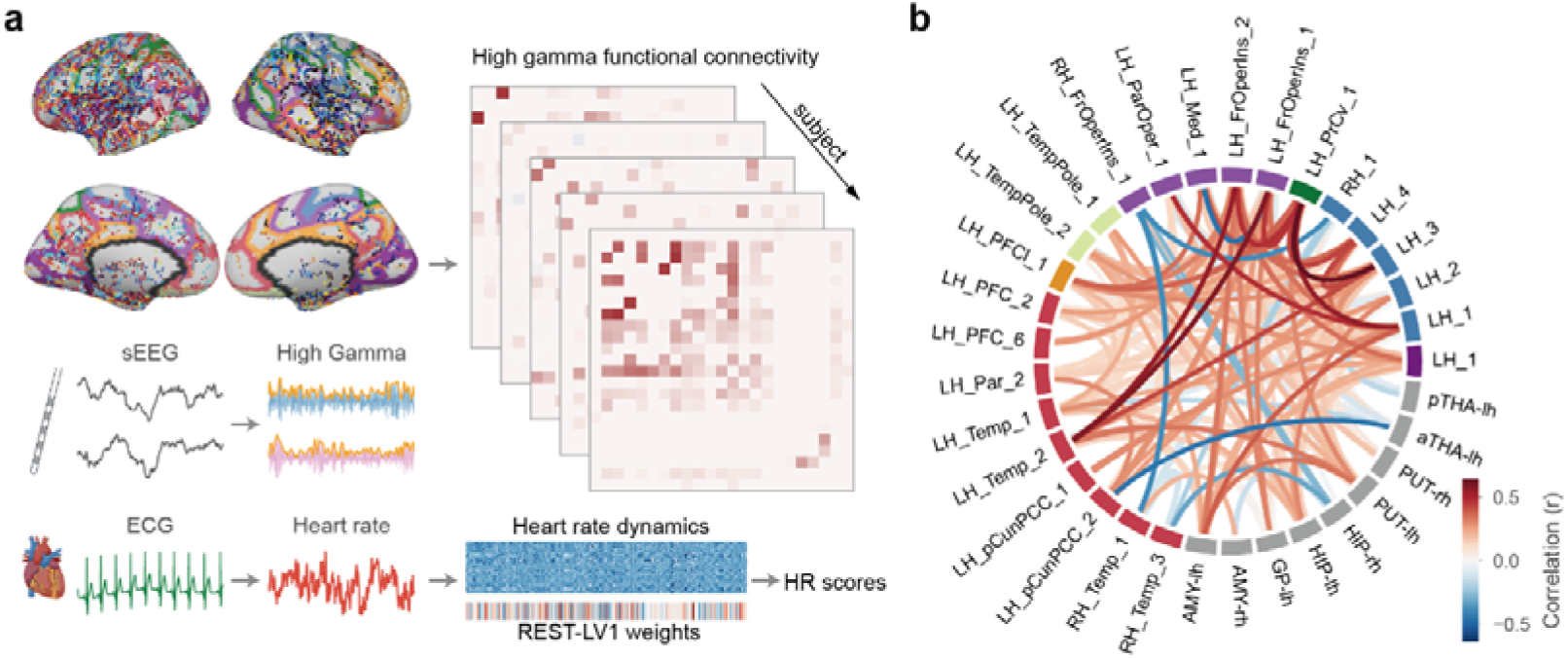
Electrophysiological validation using intracranial sEEG. **a**, Schematic workflow for neural origin validation in an independent patient cohort. Intracranial sEEG recording sites are anatomically mapped to brain regions (top left), and high-gamma band (70–120 Hz) envelopes are extracted to construct regional functional connectivity matrices across subjects (top right). Simultaneously recorded ECG signals are converted to heart rate series to extract individual heart rate dynamical properties, which are then projected onto the HCP-derived weight vector to yield subject-specific heart rate scores (bottom right). **b**, The Pearson correlation (*r*) between individual heart rate scores and sEEG-derived high-gamma functional connectivity across functional connectivity edges available in at least 10 patients. Regions are grouped by Yeo-7 network assignments, and the color of the connecting bands represents the correlation coefficient.

Together, these validations across physiological confounds control, cross-session segregation, and electrophysiological replication convergingly demonstrate that the heart-brain covariance is a stable, biologically grounded neurophysiological trait.

### Heart–brain coupling extends to active cognitive states

Building upon the robust heart-brain covariation established above, we reasoned that if this macroscopic covariance represents a core neurophysiological trait, its presence should not be confined to the spontaneous fluctuations in the resting state. To explore this, we investigated whether individual differences in task-concurrent heart rate dynamical properties reflect variations in task-evoked neural activation patterns, and how their specific loading configurations might adapt during task execution.

To ensure these peripheral profiles remain viable individual markers under cognitive load, we first evaluated whether heart rate dynamical properties maintain their individual distinctiveness across seven distinct cognitive tasks, including working memory (WM), social, emotion, language, motor, relational, and gambling. Heart rate dynamical properties across multiple scanning runs within the same task consistently exhibited greater similarity than those across different tasks (Supplementary Fig. 5a). Furthermore, the identifiability index remained consistently above threshold across all tasks (Supplementary Fig. 5b). Notably, when resting-state and task-state runs were pooled, cross-state identifiability remained robust (Supplementary Fig. 5b). Collectively, these findings demonstrate that heart rate dynamical properties comprise both state-specific and individual-specific components during task execution.

We next examined whether heart rate dynamical properties covaried with individual task activation profiles. For each subject, we extracted 17 task-evoked activation maps spanning the seven tasks (Figure 6a). Independent PLS analyses were conducted for each task to identify latent variables between heart rate dynamical properties and task-evoked activation patterns. Following a joint evaluation of the proportion of explained covariance, cross-validation generalizability, permutation-based significance, and the asynchronous cross-run projection validation (Figure 6b, c; Supplementary Fig. 6), we identified four robust latent variables: WM–LV1 (explained covariance = 44%, *p* = 0.001; training *r* = 0.24, validation *r* = 0.13), GAMBLING–LV1 (explained covariance = 59%, *p* = 0.0002; training *r* = 0.22, validation *r* = 0.16), SOCIAL–LV1 (explained covariance = 54%, *p* = 0.0002; training *r* = 0.20, validation *r* = 0.13), RELATIONAL–LV1 (explained covariance = 51%, *p* = 0.0117; training *r* = 0.21, validation *r* = 0.11).

**Figure 6.**
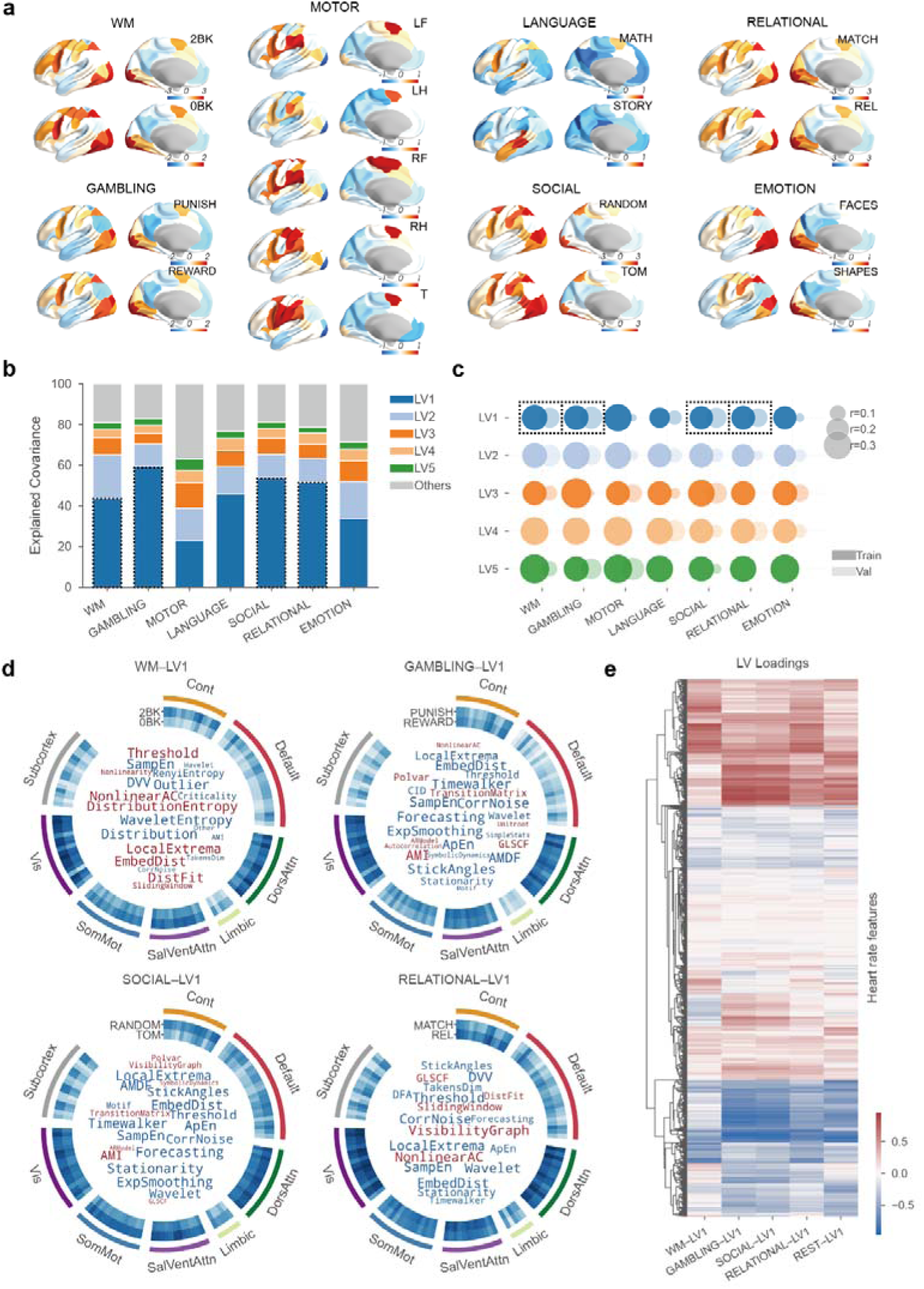
Mapping heart rate dynamical properties onto task-evoked brain activation profiles. **a**, Examples of task-evoked brain activation maps (*z*-stat) across seven cognitive tasks. **b**, Proportion of explained covariance for the top five latent variables (LV1–LV5) derived from independent PLS analyses performed for each task. **c**, Cross-validation performance across tasks for the top five latent dimensions. The size of the circles represents the mean correlation coefficients (*r*) for the training (dark shading) and test (light shading) folds across 30 repetitions. **d**, Joint profiles of brain activation and heart rate features loadings for the identified significant latent variables. The outer circular tracks illustrate the brain activation loadings grouped by Yeo-7 functional networks. The center inset word clouds present the heart rate feature loadings, with word size reflecting the absolute loading magnitude. **e,** Heart rate feature loadings across the identified task-state and resting-state latent variables. The heart rate features are ordered using hierarchical clustering to group features with similar loading profiles across different states. The color scale represents the magnitude and direction of the feature loadings.

Figure 6d illustrates the loadings of heart rate dynamical properties and corresponding task activation maps. Overall, heart rate features covarying with task-evoked activation resembled those associated with RSFC (Fig. 6e), with loading similarities ranging between 0.69 and 0.85 (Pearson’s correlation coefficient). However, while maintaining a similar pattern of negative complexity loadings and positive predictability loadings, as well as a shared focus on short timescales, loadings corresponding to specific mathematical approaches differing across task states (Supplementary Table 1). Under task conditions, predictability was strongly represented by low-order lag autocorrelation, whereas specific derived forms such as auto-mutual information (AMI) showed higher loadings in specific tasks.

For GAMBLING–LV1, higher heart rate autocorrelation was associated with negative activation loadings across both reward and punishment conditions. This association was stronger during reward conditions than punishment, and the corresponding activation patterns were predominantly distributed across visual, ventral attention, dorsal attention, and frontoparietal control networks. For RELATIONAL–LV1, higher autocorrelation is associated with negative activation loadings during both relational reasoning and match processing conditions. For SOCIAL–LV1, lower temporal complexity was associated with negative activation loadings during both random motion and Theory of Mind (TOM) conditions. In contrast, although the activation loading pattern of WM–LV1 was broadly aligned with those of the other tasks, its heart rate feature loading profile was distinct and was dominated by measures of distributional shape and non-Gaussianity.

Together, these findings indicate that heart–brain covariance extends from rest to active cognitive states while showing task-specific reconfiguration of the contributing heart rate features. Across the identified task-state latent variables, higher heart rate autocorrelation and lower temporal complexity was generally associated with lower task-evoked brain activation.

### Convergence of brain-covarying and cognitive-predictive heart rate dynamical properties

Whether the information within individual heart rate dynamics that captures RSFC and task response patterns translates into observable variations in human behavior remains a critical question. To evaluate the behavioral relevance of these heart rate signatures, we compiled each participant’s resting-state and task-concurrent heart rate scores from the brain-covarying latent variables into a 5-dimensional profile. Meanwhile, we compiled 73 behavioral traits spanning alertness, cognition, emotion, motor, personality, sensory, and mental health domains (Supplementary Table 2). Using a factor analysis model to extract robust behavioral dimensions, we extracted five distinct behavioral factors: crystallized cognition, fluid cognition, internalizing symptoms, externalizing symptoms, and social support (Supplementary Fig. 7; Supplementary Table 3).

The heart rate scores of brain-covarying latent variables are predominantly associated with crystallized and fluid cognition, whereas WM–LV1 showed weaker associations (Fig. 7a). To assess the extent to which these latent variables capture cognition-related variance in heart rate dynamics, we first trained multi-kernel ridge regression models using unprojected heart rate dynamical properties to predict crystallized and fluid cognition. Model performance was evaluated using a nested 10-fold cross-validation with 30 repetitions. We then trained equivalent models after deflating the variance explained by these brain-covarying latent variables. Removing this variance significantly reduced prediction accuracy of both crystallized and fluid cognition (Fig. 7b).

**Figure 7.**
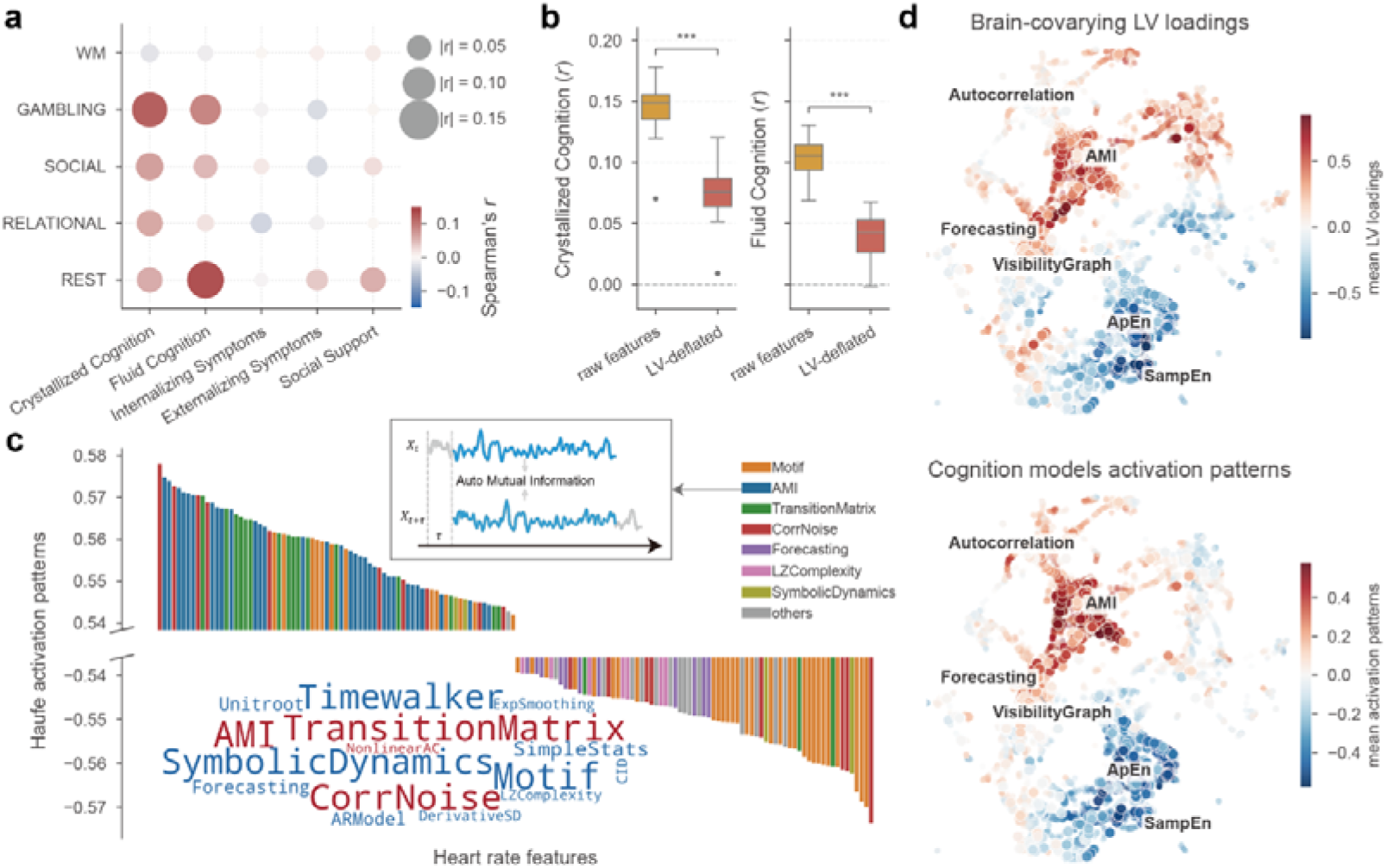
Convergence of brain-covarying and cognitive-predictive heart rate dynamical properties. **a**, Correlation matrix between the heart rate scores of brain-covarying latent variables and five distinct behavioral factors. Circle size indicates the absolute Spearman correlation coefficient (|r|), and the color scale represents the correlation magnitude and direction. **b**, Comparison of prediction accuracy (Pearson r) for crystallized and fluid cognition using multi-kernel ridge regression models trained on raw heart rate features (yellow; raw features) versus features deflated by the variance of brain-covarying latent variables (red; LV-deflated) (two-sided Mann-Whitney U tests; \*\*\**p* < 0.001). **c**, Haufe activation patterns for cognitive predictions. The top 150 positive and negative feature activations are ranked by their mean values across models and color-coded by their algorithmic classes. The word cloud displays feature classes sized proportionally to their activation magnitudes. **d**, UMAP embeddings mapping of the mean brain-covarying latent variable loadings (top) and the mean cognition model activation patterns (bottom). The color scale denotes the magnitude and direction of the loadings and activation patterns.

To characterize the heart rate dynamical properties contributing to cognitive prediction, we applied the Haufe transform^61,62^ to derive feature activation patterns. The activation patterns were highly similar across states and cognitive prediction targets (Supplementary Fig. 8a). Higher cognitive scores were associated with greater autocorrelation, lower temporal complexity, and lower forecasting errors (Fig. 7c; Supplementary Table 4). Although the predictive models were trained independently of the brain functional features, the resulting heart rate feature activation patterns showed substantial alignment with the brain-covarying loading patterns (Fig. 7d; Supplementary Fig. 8b). For instance, the GAMBLING–LV1 loading pattern was highly similar to the corresponding activation patterns for prediction of crystallized cognition (*r* = 0.87) and fluid cognition (*r* = 0.83).

Together, these findings demonstrate that neural projections distill the cognitive predictive capacity of heart rate dynamics, revealing that individual brain function and cognitive traits share a similar representation within heart rate fluctuation patterns.

## Discussion

In this study, we establish well-characterized heart rate dynamical properties as an individual-specific physiological fingerprint that maps onto large-scale neural architecture. We identify a covariance axis linking reduced temporal complexity and increased predictability of heart rate to elevated resting-state functional connectivity. We verified this heart-brain covariance as a stable neurophysiological trait through external cohort replication, regression of physiological confounds, cross-session stability testing, and direct electrophysiological confirmation. Furthermore, this coupling architecture persists across task-evoked states, and the derived brain-covarying latent variables predict cognitive performance. These findings demonstrate a convergence among peripheral autonomic dynamics, large-scale neural functional organization, and cognitive phenotypes.

Heart rate is among the most widely collected physiological signals in clinical and ambulatory settings. Rather than relying on conventional time- and frequency-domain HRV indices, we employed the hctsa framework to comprehensively capture the temporal profiles of heart rate. While this method has been successfully applied to neural signals like BOLD^63–65^ and MEG^66^, our previous work established its efficacy in characterization of heart rate dynamics during brain state transitions^31^. This approach circumvents the subjectivity and potential omissions of incremental descriptor selection while maintaining an explicit feature calculation workflow, thereby bridging high representation capacity with rigorous interpretability. Such computational transparency ensures that the key features identified in our study can be directly replicated and quantified in future research, thereby facilitating the systematic investigation of their underlying biological mechanisms.

The dynamical features identified here, including measures of entropy, autocorrelation, predictability, and visibility graph–based temporal organization, provide a broader perspective on the cardiac temporal profile. Our findings align with recent evidence demonstrating that heart rate entropy is selectively predictable from whole-brain effective connectivity^26^. However, unlike traditional HRV indices with established physiological interpretations^1,67^, the biological basis of these features remains incompletely understood. Pharmacological studies showing limited effects of vagal blockade on approximate entropy further suggest that cardiac complexity does not simply represent activity of a single autonomic pathway^68^. Instead, these properties may emerge from nonlinear interactions among sympathetic and parasympathetic regulation^41,69^, requiring further research to fully clarify the underlying physiological mechanism.

Previous studies investigating cardiovascular–brain interactions have largely focused on localized pathways, including prefrontal-amygdala, frontal-insular, and other components of the central autonomic network^10,70,71^. Our findings extend this framework by demonstrating that heart-brain coupling involves distributed functional organization encompassing default mode, salience/ventral attention, dorsal attention, and sensorimotor networks. The involvement of prefrontal and insular regions is consistent with their established roles in autonomic regulation^70,72–74^. Meanwhile, the contribution of sensorimotor regions, which are not classical CAN regions, is supported by recent evidence that optogenetic activation of motor cortex can modulate heart rate^75^. Together, our findings integrated with recent anatomical tracing and cortical stimulation evidence^12,15,76^, suggest an expansion of the functional boundaries of CAN.

The spatial relationships between the identified heart–brain covariance pattern and PET-derived cholinergic, noradrenergic, and serotonergic maps provide additional insight into the biological organization underlying this heterogeneous pattern. Cholinergic projections from the basal forebrain contribute to large-scale cortical co-fluctuation and attentional states^77^, whereas the locus coeruleus-noradrenergic system regulates arousal, attention, adaptive behavior, and autonomic responses through widespread projections^78–81^. Serotonergic systems have also been implicated in coordinating behavioral state and cardiovascular regulation^75,82^. Together, these neuromodulatory systems represent potential biological contributors linking brain-wide functional states with autonomic regulation. However, how inter-individual variations within these neuromodulatory systems shape individual differences in whole-brain dynamics remains largely unexplored, primarily due to the historical scarcity of scalable detection techniques suitable for large-scale cohorts to precisely quantify individual neuromodulatory profiles. Furthermore, whether and how these central neuromodulatory centers govern the autonomic output to drive downstream heart rate dynamics or are conversely modulated by feedback from peripheral autonomic afferents, requires further elucidation.

Despite the statistical robustness of the observed heart-brain covariance and its spatial alignment with neuromodulatory gradients, a critical challenge in studying cardiovascular–brain interaction is distinguishing biological coupling from shared physiological confounds. Both cardiac signals and neuroimaging measures can be influenced by non-neural physiological fluctuations^57^ or environmental perturbations^57,83^. The persistence of the identified covariance after accounting for these potential confounding factors supports the interpretation that brain-heart covariance reflects a stable brain-body shared trait. The convergence between fMRI-derived and sEEG-derived patterns further suggests a conserved neurogenic mechanism that transcends macro-scale hemodynamic limitations.

Building upon this robust intrinsic covariance, a natural next question is whether this heart-brain coupling persists under task-evoked states or is homogenized by shared task constraints. While previous studies established that cognitive load generally drives cardiac rhythms into higher-complexity states at the group level^20,84^, our findings extend this notion by demonstrating that individual variations in these task-bound heart rate dynamical properties index the degree of personalized brain activation. This relationship is consistent with the neurovisceral integration model, in which higher-order cortical systems contribute to autonomic regulation during adaptive behavior^27,85^. At the same time, accumulating evidence supports bidirectional interactions through interoceptive pathways, suggesting that cardiac dynamics may reflect reciprocal communication between central and peripheral systems^85,86^. Therefore, the exact causal mechanisms underlying this association remain largely elusive. The preservation of individual identification capabilities, along with the presence of similar covariance patterns during cognitive tasks, reflects a blend of intrinsic neurobiology and analytical constraints. Neurobiologically, this high convergence aligns with the consensus that intrinsic and task-evoked brain organizations share a substantial generalized scaffold^87–89^. Computationally, time series profiling incorporates non-execution epochs may blunt task-specific variance. Against this backdrop, dissecting the precise physiological substrates that underwrite these mathematically convergent or divergent features becomes critical. Future interventional studies leveraging similar comprehensive characterization will be essential to assign explicit biological annotations to these mathematical metrics.

Extracting behaviorally relevant representations grounded in robust neurophysiological foundations from accessible peripheral signals offers a scalable strategy for biomarker development. We found that projecting heart rate dynamical properties onto latent dimensions that covary with neural features simultaneously captures variance relevant to crystallized and fluid intelligence, consistently echoing the premium predictive capacity of neuroimaging features for cognitive traits^62,90^. Specifically, elevated autocorrelation and reduced heart rate complexity correlated with higher cognitive performance. At the neural level, the lower task-evoked brain activation accompanying this axis aligns with the neural efficiency hypothesis, which posits that higher cognitive capacity requires less neural resource consumption^91–94^. The pattern parallels evidence from blood pressure signals, where higher entropy is similarly linked to lower cognitive performance^95^. However, to prevent semantic confusion and the misuse of broad terminology, it must be emphasized that lower short-term entropy does not equate to lower overall sequence variability. In fact, these two features exhibit a negative correlation in our dataset. As the key features identified here operate on time scales fundamentally distinct from those of classical HRV indices^96^, these findings underscore the necessity of exploring the distinct physiological implications and methodological specificities of heart rate features across different time scales.

The clinical and translational relevance of these findings lies in the accessibility of cardiovascular signals. ECG and PPG are routinely collected in clinical, ambulatory, and wearable settings, enabling longitudinal characterization at a scale that is difficult to achieve with neuroimaging. Comprehensive characterization of cardiac signals may therefore provide a scalable framework for studying cardiovascular–brain interactions across aging, cardiovascular disease, and neurological conditions. Importantly, we provide a theoretical rationale for deriving physiological biomarkers from heart rate dynamics, suggesting that their predictive utility is anchored in shared information with the central nervous system rather than in opaque, black-box associations. However, we emphasize that the current findings represent a proof-of-concept rather than an established biomarker. Given the modest effect sizes observed both in our study and in prior research^97^, and echoing the lessons from brain-wide association studies^98^, establishing viable physiological biomarkers will demand much larger-scale, multi-center validation, an endeavor made highly feasible by the ubiquitous nature of heart rate monitoring.

It is worth emphasizing that the primary aim of this study was to establish the presence of a coordinated heart–brain–behavior pattern and provide insights for future mechanistic investigations, rather than to infer causality. Decoupling true causal directions remains considerably more complex due to bidirectional interactions and the spatiotemporal resolution limits of current recording techniques, thereby presenting profound interpretive challenges for purely statistical causal frameworks prior to these foundational breakthroughs. Acknowledging this scope, several specific limitations of the current work warrant mention.

First, our sample was restricted to healthy young adults. Although the observed heart–brain covariation generalized to an independent cohort, subtler or condition-specific patterns may have been overlooked. Second, the neuromodulatory analyses were based on population-average positron emission tomography maps rather than individual molecular measurements and therefore provide spatial context rather than direct evidence of neurotransmitter involvement. Third, we used sEEG data from patients with epilepsy to support the neural basis of the heart–brain covariation. In addition to the limited sample size and spatial coverage, this assumes that the coupling is preserved despite the pathology, which warrants caution and further validation. Finally, we adopted a conservative strategy in LV selection, particularly for task-evoked states where independent external validation was unavailable; while this approach maximizes reliability, it may inadvertently overlook other subtle modes of heart–brain covariance. In summary, we demonstrate that well-characterized heart rate dynamics hold potential as a non-invasive window into individual neural architecture and cognitive traits. Future efforts should focus on assigning biological annotations to these mathematical features, ultimately enabling the development of biomarkers grounded in reliable neurobiological mechanisms across broader population scales.

## Methods

### Participants

We used data from the Human Connectome Project Young Adult (HCP-YA) S1200 release^39^, which comprises multimodal neuroimaging and behavioral assessments from 1,206 healthy young adults aged 22–37 years. Each participant received four resting-state fMRI (REST) scans. Each REST run lasted 14.4 minutes. In addition, participants completed seven task-based fMRI paradigms designed to probe distinct cognitive domains: working memory (∼10 minutes), language processing (∼7.5 minutes), relational processing (∼5.4 minutes), motor execution (∼7.6 minutes), emotional processing (∼2.3 minutes), social cognition (∼3.6 minutes), and gambling (∼4.2 minutes). Task designs and timing followed the standardized HCP protocol.

Physiological signals were synchronously recorded during all fMRI runs. Each participant also completed a comprehensive behavioral battery assessing cognitive ability and self-report measures.

The external validation data were from the MPI Leipzig Mind-Brain-Body (LEMON) dataset ^50^, which includes multimodal neuroimaging and physiological data from 228 healthy adults aged 20–75 years. Participants underwent resting-state fMRI acquisition and simultaneous physiological recordings.

### Imaging data acquisition and preprocessing

For HCP-YA dataset, all imaging data were acquired on a customized Siemens 3T scanner using a multiband echo-planar imaging sequence as part of the HCP protocol (TR = 720 ms, TE = 33.1 ms, flip angle = 52°, voxel size = 2.0 mm isotropic). Comprehensive acquisition parameters have been reported in the original paper^39^. We used the minimally preprocessed and ICA-FIX denoised functional data^99^. The preprocessing pipeline includes gradient distortion correction, motion correction, field map–based EPI distortion correction, alignment to structural images, normalization to MNI space, and projection to the standard grayordinate space. Spatial ICA followed by automated component classification and removal (ICA-FIX) was used to further denoise the data. In addition, global signal regression (GSR) was performed by regressing out the average time series across all cortical vertices. Both GSR and non-GSR versions of the data were retained for downstream analyses.

For the LEMON dataset, resting-state fMRI was acquired on a Siemens 3 T Verio scanner with the following parameters: TR = 1400 ms, TE = 39.4 ms, flip angle = 69°, voxel size = 2.3 mm isotropic, resulting in 657 volumes per run (∼15 min 30 s per run). The preprocessing for LEMON dataset was performed using DeepPrep^100^ (v25.1.0), including projection to the standard grayordinate space. Preprocessing reports were visually inspected, and subjects with evident quality issues were excluded from further analysis. From the preprocessed time series, linear trends were removed, and nuisance variables were regressed out, including white matter and cerebrospinal fluid signals, as well as the motion parameters and their first- and second-order temporal derivatives.

### PPG acquisition and preprocessing

For the HCP-YA dataset, physiological signals were concurrently recorded during all fMRI runs via a PPG sensor sampled at 400 Hz. For LEMON dataset, PPG signals were recorded with a sampling frequency of 1,000 Hz. PPG signals preprocessing was performed using the NeuroKit2 (v0.2.10)^101^. Raw PPG signals were bandpass filtered (0.5–8 Hz), and cardiac systolic peaks were initially detected using Elgendi’s peak detection algorithm^102^. Signal quality was then evaluated using the template-matching method, excluding runs with an average quality score below 0.90. For the retained runs, artifacts and missing peaks were corrected using Kubios algorithm^103^. Inter-beat intervals were then resampled to 4 Hz to yield uniformly sampled heart rate for subsequent time-series analysis. Each resulting heart rate series was then visually inspected and runs with prominent noise or flat signal were manually excluded.

### Functional connectivity estimation

Resting state functional connectivity (RSFC) was estimated separately for each fMRI run. Time series were extracted from 116 brain regions comprising 100 cortical parcels from the Schaefer100 atlas and 16 subcortical regions^47,48^. For each run, pairwise Pearson correlation coefficients were computed between all regional time series, yielding a 116 × 116 functional connectivity matrix. To assess the robustness of our findings to parcellation granularity, additional analyses were conducted using the Schaefer atlas with 200 and 400 cortical parcels, Glasser atlas with 360 cortical parcels^104^, combined with the same 16 subcortical regions^47^.

### Task-state fMRI activation mapping

Task-state fMRI activation maps were estimated separately for each individual task run. First-level general linear model analyses were performed on the preprocessed grayordinate time series using the Nilearn (v0.10.2) packages in Python. For each run, a design matrix was constructed by convolving the task design with a canonical Glover hemodynamic response function. To control motion-related confounds, head motion parameters and their first-order temporal derivatives, were included in the design matrix as nuisance regressors. Low-frequency drift was removed by applying a high-pass filter with a 200 s cutoff implemented via a cosine drift model. Temporal autocorrelation in the fMRI noise was modeled and corrected using a first-order autoregressive structure during model estimation. Linear contrasts of interest were computed to generate subject-level contrast parameter estimate maps and standardized *z*-stat maps for each run. Finally, the resulting dense activation maps were parceled into the same 116-region atlas used for resting-state functional connectivity analysis.

### Heart rate feature extraction

Heart rate time series were characterized by a unified, high-dimensional profile that spans both classical HRV measures and advanced dynamical properties. For each fMRI run, we first extracted conventional metrics using the NeuroKit2 (v0.2.10): SDNN, RMSSD, pNN50, low-frequency (LF) power, high-frequency (HF) power, the LF/HF ratio, and the Poincaré plot descriptor SD2. To capture a broader, more comprehensive array of mathematical properties, these conventional measures were integrated with the highly comparative time-series analysis (hctsa) framework^37,38^. The hctsa toolbox computes a comprehensive set of time-series features that capture a wide range of dynamical properties. Features that were non-informative such as those reflecting only the length of the input sequence, or returned undefined values were excluded. In addition, features exhibiting extremely low variability across individuals, defined as having interquartile range < 0.001, were removed. To facilitate comparison, only features that met these quality control criteria in both resting-state and task-based data were retained. After this filtering procedure, 6,817 features were retained and robust-sigmoid normalized for downstream analyses. Detailed algorithms for the key features identified in subsequent analyses are provided in the Supplementary Methods.

### Identifiability index

Fingerprinting of individual differences in heart rate dynamical properties was quantified using the identifiability index^42–44^, which has previously been applied to assess individual specificity in brain connectome. For the resting-state, participants with four high-quality heart rate series that passed quality control were included. For the seven task conditions, participants with two high-quality heart rate series per task were retained.

To assess the identifiability index for each acquisition paradigm, we computed the mean within-participant correlation (*μ_intra_*) and the mean between-participant correlation (*μ_inter_*) across corresponding hctsa-derived feature vectors. For cross-state identifiability, all heart rate recordings from the same participant were pooled across all conditions, resulting in 18 hctsa feature vectors per subject. The identifiability index was calculated as:

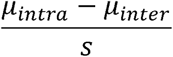

where *s* is the pooled standard deviation, yielding a measure analogous to an effect size.

Temporal profile similarity between runs was defined as the Pearson correlation between pairs of hctsa feature vectors from the same individual^64^. The average of all pairwise within-subject correlations was computed, transformed using Fisher’s *z*, averaged, and then converted back to the *r* metric for interpretation.

### PLS analysis

To identify multivariate associations between heart rate dynamical properties and brain function, we applied partial least squares (PLS) analysis separately under resting-state and task-state conditions. Compared with alternative multivariate methods, PLS offers greater stability and robustness in high-dimensional settings, particularly when the number of features exceeds the number of subjects and strong internal correlations are present^51,105^. PLS extracts orthogonal latent variables maximizing the shared covariance between two data matrices, namely a brain metrics matrix *X* of size *n* by *g* and a physiological features matrix *Y* of size *n* by *t*, where *n* denotes the number of participants, *g* denotes the number of brain features, and *t* denotes the number of heart rate dynamical properties.

In the resting-state analysis, the functional connectivity vector for each subject was defined as the Fisher’s z-transformed upper triangle of the connectivity matrix. In the task-state analysis, brain activation matrices were constructed by concatenating the 116 regional values across multiple task contrasts of interest for each task, yielding 232 features for tasks with 2 contrasts of interest (working memory, gambling, language, social, relational, and emotion) and 580 features for the motor task (which had 5 contrasts of interest). Prior to PLS, age, sex, and mean framewise displacement were regressed out from both matrices to control for potential confounds. Following confound regression, variables were standardized across subjects.

PLS was performed via singular value decomposition (SVD) on the cross-covariance matrix *C* between *X* and *Y*, defined as:

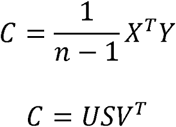

where *U* and *V* are orthonormal matrices containing the left and right singular vectors (weight vectors), and *S* is a diagonal matrix containing the singular values. The singular values reflect the covariance captured by each latent variable (LV), and the relative proportion of explained covariance for each LV was calculated as the ratio of its squared singular value to the sum of all squared singular values. Subject-level brain scores (*Score_X_*) and heart rate scores (*Score_Y_*) for each latent variable were computed by projecting the standardized data onto their respective weight vectors:

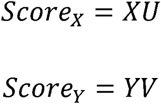

These scores quantify the individual expression of the identified brain and physiological patterns. Variable contributions were indexed by loadings, calculated as the Pearson correlation coefficient between individual variables and their corresponding subject-level scores^55^.

To evaluate the generalizability of the latent variables and prevent data leakage between related individuals, we conducted family-controlled cross-validation. For both resting-state and task-state PLS, we performed a repeated 10-fold cross-validation (repeated 30 times). The splits were grouped by family to ensure that participants belonging to the same family were kept within the same fold together and never split between training and test sets. In each split, PLS weights were derived from the training set, and scores for the held-out test set were computed by projection. Generalizability was quantified as the Pearson correlation coefficient between the projected brain and physiological scores in the test set.

To assess the statistical significance of the latent variables, we performed 10,000 permutation tests by shuffling subject labels in the physiological matrix. To avoid inflated significance of the dominant component, we did not employ the Procrustes rotation framework (where permuted singular vectors are aligned to empirical vectors), as recent methodological research has shown that rotated permutations systematically bias the null distribution and increase false-positive rates for the first latent variable^49^. We used a sequential, unrotated tail sum of squares (SSQ) permutation test^106^. For each component k, the test statistic was defined as the sum of squared singular values (SSQ) from component k to the maximum rank:

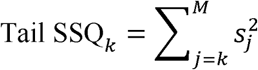

For each permutation, the tail SSQ was computed in the same manner using the permuted singular values. The empirical p-value for each component was calculated as the proportion of permutations where the permuted tail SSQ exceeded the observed empirical tail SSQ. These p-values were evaluated sequentially; component k was considered significant only if all preceding components (from 1 to k-1) met the significance threshold of *p* < 0.05. If a component failed to meet this threshold, the sequential evaluation was terminated, and all subsequent components were deemed non-significant.

To validate the stability and neurobiological validity of our resting-state PLS findings, we performed three complementary validation analyses. First, to assess sensitivity to parcellation granularity, we repeated the PLS analysis using functional connectivity matrices constructed with 216-region, 376-region, and 416-region cortical parcellations. Second, to examine whether the heart-brain coupling was driven by global physiological fluctuations in the fMRI BOLD signal, we repeated the PLS analysis using global signal regression (GSR) preprocessed connectivity matrices and also projected the GSR-preprocessed data directly onto the weight vectors derived from the non-GSR PLS model. Third, to determine whether the identified covariance represents a stable, trait-like signature rather than transient, state-dependent synchronization during a single session, we performed a cross-session projection analysis. Using data from participants with multiple resting-state sessions, we constructed two cross-paired datasets by replacing one modality of the data with that of a non-concurrent run before projecting the data matrices onto the original PLS weight vectors to obtain cross-paired brain and heart rate scores. The stability and significance of this cross-paired coupling were evaluated by computing the Pearson correlation coefficient between the projected scores across participants.

### Respiration-related confounds control analysis

To evaluate whether the multivariate association between brain functional connectivity and heart rate dynamical properties was driven by respiration-related physiological confounds, we performed a covariate control analysis using concurrent respiration recordings. For each participant, we extracted five key metrics using the NeuroKit2 (v0.2.10): Porges-Bohrer respiratory sinus arrhythmia (RSA), mean respiration rate, respiratory volume per time, mean inspiration duration, and the inspiration/expiration duration ratio. We then regressed these five metrics out of both RSFC scores and heart rate scores using ordinary least squares linear regression. Finally, Pearson correlation coefficients were computed between the residual brain and physiological scores to test the statistical significance of the coupling after controlling for respiration-related confounds.

### Annotation with biological maps

We obtained 20 brain maps from the neuromaps toolbox (v0.0.5)^54^, encompassing previously reported or curated data on neurotransmitter system distribution^55^. All maps were parcellated into the Schaefer-100 atlas using the corresponding spatial reference. To assess the spatial similarity between regional importance and each cortical map, we computed Spearman correlation coefficients. Given the spatial autocorrelation inherent in cortical data, we performed spatial autocorrelation-preserving permutation test^56^, repeating the procedure 10,000 times to generate null distributions. Resulting p-values were corrected for multiple comparisons using the FDR procedure, with FDR < 0.05 considered statistically significant.

### Intracranial stereo-electroencephalography acquisition and analysis

Intracranial stereo-electroencephalography (sEEG) and concurrent ECG data were collected from 41 patients undergoing presurgical evaluation for epilepsy localization at Huashan Hospital. The study protocol was approved by the Ethics Committee of Huashan Hospital (KY2019-518). Continuous signals were recorded using a digital Nihon Kohden system at a sampling rate of 2,000 Hz. Resting-state epochs were visually inspected and selected by an experienced clinical neurologist to ensure they were completely free of epileptiform activity and artifacts. The duration of the extracted time segments was kept consistent with the acquisition length of the REST protocol in HCP-YA dataset. The sEEG signals were bipolar re-referenced and bandpass filtered between 0.5 and 250 Hz, with a notch filter applied to remove powerline noise. The data were then filtered into the high-gamma band (70–120 Hz) using a finite impulse response filter. Amplitude envelopes were extracted via the Hilbert transform and subsequently down sampled to 40 Hz. Functional connectivity was evaluated using pairwise Pearson correlations of these envelopes. For anatomical localization, sEEG electrode contacts were co-registered to MNI space. The geometric midpoint of each bipolar channel was mapped to the 116-node atlas. Heart rate time series were extracted from the ECG data, and features were extracted using the same framework. These features were then projected onto the HCP-derived resting-state latent variable weight vector to obtain individual heart rate scores. Due to the sparse spatial coverage of sEEG, cross-subject analysis was restricted to functional connectivity edges present in at least 10 patients, resulting in a final set of 153 edges. For each edge, the relationship between connectivity strength and individual heart rate scores was evaluated using Pearson correlation.

### Behavioral prediction

To reduce the dimensionality and extract robust behavioral traits, we performed a factor analysis on 73 behavioral items, spanning alertness, cognition, emotion, motor, personality, sensory, and mental health domains. The factor analysis was performed using a varimax rotation, extracting five behavior factors: crystallized cognition, fluid cognition, internalizing symptoms, externalizing symptoms, and social support.

To evaluate the direct behavioral relevance of the brain-covarying physiological signatures, we aligned each participant’s 5-dimensional latent heart rate profile (comprising the heart rate score from REST and the four task states: WM, GAMBLING, SOCIAL, and RELATIONAL) with their behavior factor scores, resulting in a final cohort of 561 participants with complete records across all modalities. Then we computed Spearman correlation coefficient between the heart rate scores and the behavior factor scores.

To establish a baseline and verify whether the brain-covarying latent components efficiently capture the core behaviorally relevant variance, we trained a multi-kernel ridge regression (MKRR) model directly on the raw, unprojected high-dimensional features, integrating physiological information across all five states. Following standardization, a state-specific linear kernel matrix was computed for each of the five states, representing the pairwise similarity of heart rate temporal profiles between participants within that specific context. These five resulting kernel matrices were then stacked to construct a composite multi-kernel training matrix. We solved the MKRR model to predict the five behavior factors using the random search solver implemented in the himalaya (v0.4.11) package. This solver jointly optimizes both the kernel weights and the global regularization parameter via an inner 10-fold cross-validation loop. To evaluate whether the behavioral predictions achieved by the heart rate dynamical properties were mainly driven by the brain-covarying PLS latent variables, we also trained a MKRR model on the raw features after deflating the latent variables in the kernel space. Within each cross-validation fold, the state-specific raw kernel matrices (*K*_raw_) were orthogonalized with respect to the latent scores (*P*) using Gram-matrix orthogonalization:

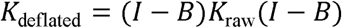

where *B* = *P*(*P^T^P*)^−1^*P^T^* represents the projection matrix onto the latent space. This deflation was computed on the training set and applied out-of-sample to the test set. The deflated kernel matrices across all states were then stacked and solved using the same MKRR framework.

To prevent familial dependencies in the HCP-YA cohort from inflating prediction performance, a family-controlled splitting strategy was enforced across all models. In both outer and inner cross-validation loops, participants from the same family were always assigned to the same fold together, ensuring that family members were never split between the training and test sets. Prediction accuracy was quantified using the Spearman correlation coefficient (*r*) between the predicted and observed behavioral factor scores. For each of the 30 repetitions, the correlation coefficients across the 10 folds were Fisher’s *z*-transformed, averaged, and then back-transformed to the Spearman *r* metric. The final prediction performance was represented as the mean correlation coefficient across the 30 repeats. Model comparisons across the 30 repeats were evaluated using two-sided Mann-Whitney U tests.

To interpret the contributions of heart rate features to the behavioral predictions, we applied the Haufe transform to convert the backward decoding regression coefficients from the raw-feature-based MKRR models into forward activation patterns^61^. To map the predictive outputs back to the original feature space, we computed a feature-level activation pattern vector:

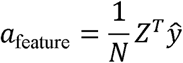

where *Z* is the feature matrix, *ŷ* is the prediction output. This formulation yields a 6,817-dimensional activation pattern vector for each state, representing how specific heart rate dynamical properties relate to behavioral predictions.

## Data availability

The Human Connectome Project Young Adult (HCP-YA) dataset is publicly available via ConnectomeDB (https://balsa.wustl.edu/). The MPI Leipzig Mind-Brain-Body dataset is publicly accessible via OpenNeuro under accession code ds000221 (https://openneuro.org/datasets/ds000221). The sEEG dataset can be provided by the corresponding authors upon reasonable request for academic research purposes.

## Code availability

Code for analyses is available via GitHub at https://github.com/Qian-Liyi/HBDimension.

## Acknowledgments

We thank the Human Connectome Project for providing neuroimaging and physiological data. We express our gratitude to all the patients and their families for their participation and cooperation in this study.

## Author contributions

L.Y.Q. contributed to conceptualization, methodology, formal analysis, and wrote the original draft of the manuscript. Z.Y.X. and B.D.F. contributed to methodology and writing – review & editing. S.H.J. and Y.M.X. contributed to conceptualization and writing – review & editing. Y.H.X. was responsible for data curation. L.C. contributed to conceptualization, funding acquisition and writing – review & editing. G.S. contributed to conceptualization and writing – review & editing. Y.Q.Z. and Y.Z.Z. contributed to writing – review & editing. X.Z., K.S., and Y.M. contributed to conceptualization, supervision, funding acquisition, project administration, and writing – review & editing.

## Sources of funding

This work was supported by National Science and Technology Major Project (2025ZD0215100 to L.C.).

Brain Science and Brain-like Intelligence Technology-National Science and Technology Major Project (2025ZD0219100 to K.S.)

Shanghai Municipal Health Commission Collaborative Innovation Group (2024CXJQ03 to Y.M.)

Science and Technology Innovation Plan of Shanghai Science and Technology Commission (23Y31900300 to Y.M.)

National Natural Science Foundation of China (82272116 to L.C., 82472244 to X.Z., and 82301642 to K.S.)

## Disclosures

Authors declare that they have no competing interests.

## Supplemental materials

Supplementary Methods Supplementary Figures 1–8

Supplementary Tables 1–4

## Notes

### Competing Interest Statement

The authors have declared no competing interest.

