## Supplementary methods and figures for "Heart rate dynamics embed a shared representation of individual brain organization and cognitive function"

### Heart rate dynamics embed a shared representation of individual neural architecture and cognitive traits

#### Supplementary Methods

##### Methods for key features

Approximate Entropy (ApEn) measures the complexity of a time series $y$ of length $N$ by calculating the probability that patterns of length $m$ that are close within a similarity threshold $r$ remain close when the pattern length is extended to $m+1$. For $N-m+1$ reconstructed vectors $x_{i}=\left[ y_{i} , y_{i+1} , \ldots, y_{i+m-1} \right]$, the matching probability is:

$$C_{i}^{m}\left( r \right)=\left( N-m+1 \right)^{-1}\sum_{j=1}^{N-m+1} \Theta\left( r-d\left( x_{i} , x_{j} \right) \right)$$

where $d\left( \cdot, \cdot\right)$ is the Chebyshev distance, and $\Theta$ is the Heaviside step function. $ApEn$ is defined as:

$$\text{ApEn}\left( m , r \right)=\Phi^{m}\left( r \right)-\Phi^{m+1}\left( r \right)$$

where the average log-probability is:

$$\Phi^{m}\left( r \right)=\left( N-m+1 \right)^{-1}\sum_{i=1}^{N-m+1} \ln\left( C_{i}^{m}\left( r \right) \right)$$

with default parameters $m=1$ and $r=0.2\sigma$ (where $\sigma$ is the standard deviation of the time series).

Sample Entropy (SampEn) eliminates the self-matching bias of ApEn by excluding self-matches ($j\neq i$), defined as:

$$\text{SampEn}\left( m, r \right)=-\ln\left( A^{m}\left( r \right)/B^{m}\left( r \right) \right)$$

where $B^{m}\left( r \right)$ and $A^{m}\left( r \right)$ are the matching probabilities for vectors of length $m$ and $m+1$, respectively. Default parameters are $m=2$ and $r=0.1\sigma$. We also computed the Quadratic Sample Entropy, defined as $SampEn+\ln\left( 2r \right)$.

Multiscale Entropy (MSE) measures complexity across multiple timescales. The time series is coarse-grained at scale factor $\tau$ by averaging data points within non-overlapping windows:

$$y_{j}^{\left( \tau\right)}=\frac{1}{\tau}\sum_{i=\left( j-1 \right)\tau+1}^{j\tau} y_{i}, 1\leq j\leq\left\lfloor N/\tau\right\rfloor$$

and $SampEn$ ($m=2,r=0.15\sigma$) is computed for each coarse-grained series $y^{\left( \tau\right)}$ across scales $\tau\in\left[ 1 , 10 \right]$.

Visibility Graph analysis maps the time series to a complex network. We constructed Horizontal Visibility Graphs (HVGs) where each time point is a node, and nodes $i$ (at time $t_{i}$) and $j$ (at time $t_{j}$) are connected if all intermediate values $y_{k}$ lie below the horizontal line connecting them:

$$y_{k}<\min\left( y_{i} , y_{j} \right), \forall t_{k}\text{ s.t. }t_{i}<t_{k}<t_{j}$$

Time series longer than 5000 points were truncated to their first 5000 samples to ensure tractability. Various topological features were extracted from the degree distribution $k$, including its moments, entropy, and goodness-of-fit to Gaussian, exponential, and power-law distributions.

Local Simple Fit evaluates short-term predictability of the time series. For a training length $L$ (default $L=3$), the next value $y_{t+1}$ is predicted using the local mean:

$$\hat{y}_{t+1}=\frac{1}{L}\sum_{i=0}^{L-1} y_{t-i}$$

or using the local median:

$$\hat{y}_{t+1}=\text{median}\left( y_{t-L+1} , \ldots, y_{t} \right)$$

or using the local linear fit:

$$\hat{y}_{t+1}=\beta_{0}+\beta_{1}\left( L+1 \right)$$

where $\beta_{0},\beta_{1}$ are estimated from ordinary least squares regression over the past $L$ values. The prediction errors $e_{t}=\hat{y}_{t}-y_{t}$ were analyzed to compute features reflecting forecast bias, error variance, sliding-window stationarity of errors, and residual autocorrelation.

Linear Autocorrelation measures linear dependencies at time lag $\tau$. The normalized autocorrelation function (ACF) $\rho\left( \tau\right)$ is computed via the Fast Fourier Transform (FFT) leveraging the Wiener-Khinchin theorem:

$$\rho\left( \tau\right)=\frac{\mathcal{F}^{-1}\left\{ \mathcal{\mid F}\left\{ y-\mu\right\}\mid^{2} \right\}}{\sigma^{2}}$$

where $F$ is the Fourier Transform, $\mu$ is the mean of the time series, and $\sigma^{2}$ is the variance. This provides an efficient calculation of linear correlations across all lags.

Nonlinear Autocorrelation captures higher-order statistical dependencies. These features are formulated as joint expectations of the z-scored time series $x$ across a vector of time delays $\tau=\left[ \tau_{1} , \tau_{2} , \ldots, \tau_{m} \right]$:

$$\text{NLAC}\left( \tau\right)=\left\langle x_{t}\cdot x_{t-\tau_{1}}\cdot x_{t-\tau_{2}}\cdot\cdots\cdot x_{t-\tau_{m}} \right\rangle$$

For configurations involving an odd number of variables, the mean of the absolute product was taken to avoid cancellations from sign fluctuations:

$$\text{NLAC}_{\text{abs}}\left( \tau\right)=\left\langle\mid x_{t}\cdot x_{t-\tau_{1}}\cdot x_{t-\tau_{2}}\cdot\cdots\cdot x_{t-\tau_{m}}\mid\right\rangle$$

Automutual Information (AMI) measures both linear and non-linear shared information between the time series and its lagged version $y_{t-\tau}$. The mutual information $I\left( y_{t} ; y_{t-\tau} \right)$ is defined as:

$$I\left( y_{t} ; y_{t-\tau} \right)=\iint p\left( y_{t} , y_{t-\tau} \right)\ln\left( \frac{p\left( y_{t} , y_{t-\tau} \right)}{p\left( y_{t} \right)p\left( y_{t-\tau} \right)} \right)dy_{t}dy_{t-\tau}$$

Non-parametric AMI estimation was performed using the Kraskov-Stögbauer-Grassberger (KSG) nearest-neighbor entropy estimators (specifically Algorithm 1 and 2) with $k=3$ neighbors and zero added noise from the Java Information Dynamics Toolkit (JIDT). A parametric Gaussian estimator, solved analytically as:

$$I\left( y_{t} ; y_{t-\tau} \right)=-0.5\ln\left( 1-r^{2} \right)$$

 where $r$ is the Pearson correlation coefficient, was also computed for comparison.

#### Supplementary Figures

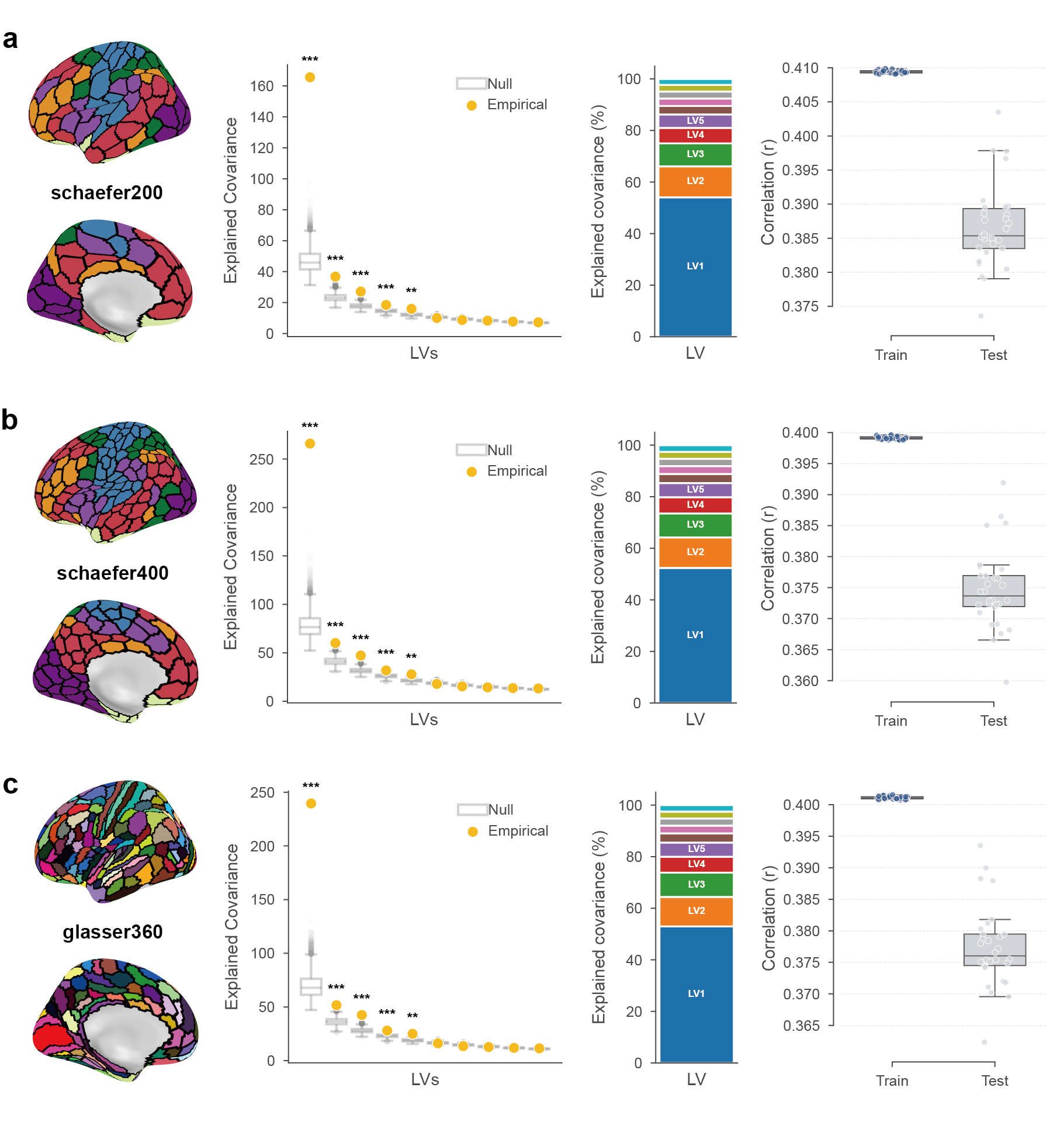

**Figure S1 | Robustness of resting-state heart-brain covariance across varying spatial parcellation resolutions. a–c**, Evaluation of the PLS model generalizability and significance using functional connectivity matrices constructed with cortical parcellations of varying spatial resolutions: 216-region (**a**), 416-region (**b**), and 376-region (**c**) atlases. Each panel presents the explained covariance of LVs against 10,000 permutation nulls (left; **p* < 0.05, ***p* < 0.01, ****p* < 0.001), the relative proportion of explained covariance (middle), and the cross-validation performance (right).

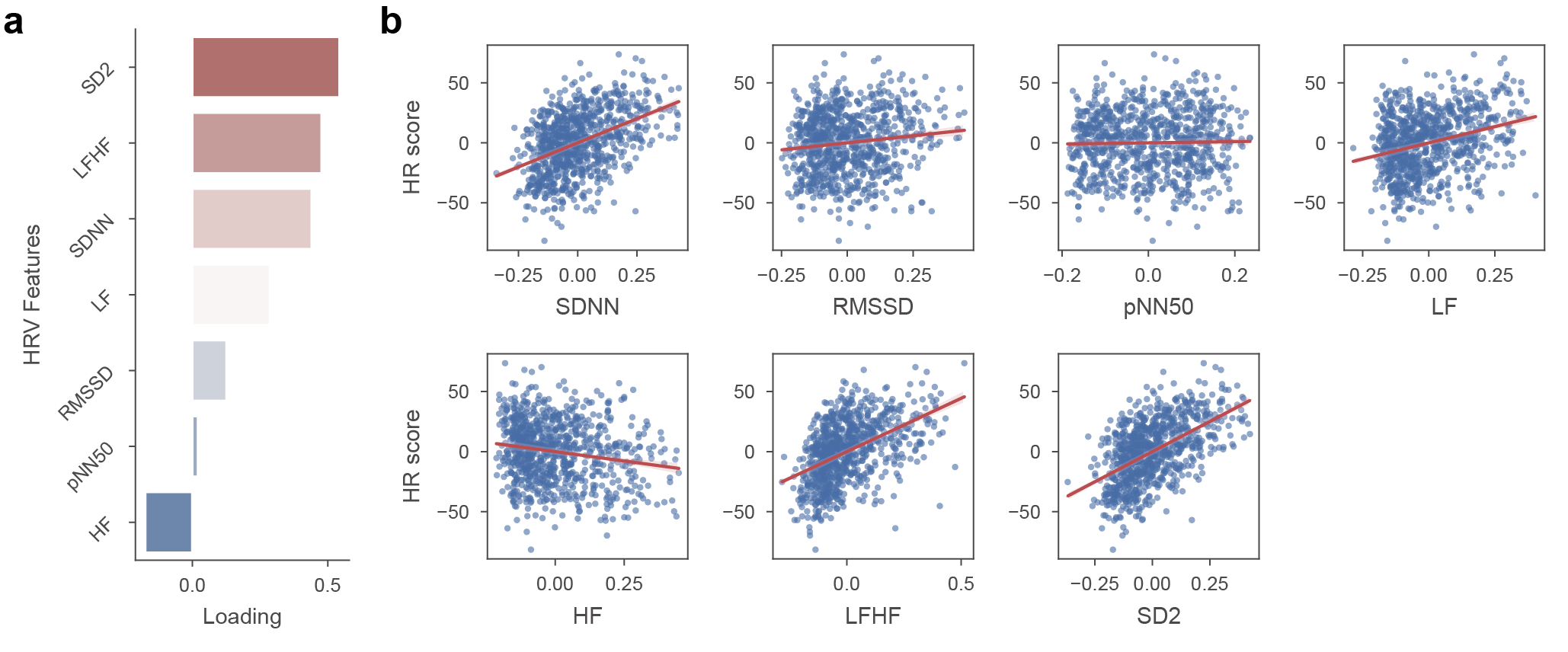

**Figure S2 | Contribution of classic heart rate variability metrics to the primary latent variable in resting-state. a,** The PLS loadings of standard time-domain and frequency-domain HRV features on the primary latent variable covarying with resting-state functional connectivity. **b,** The relationships between the individual HR scores and these specific conventional HRV metrics.

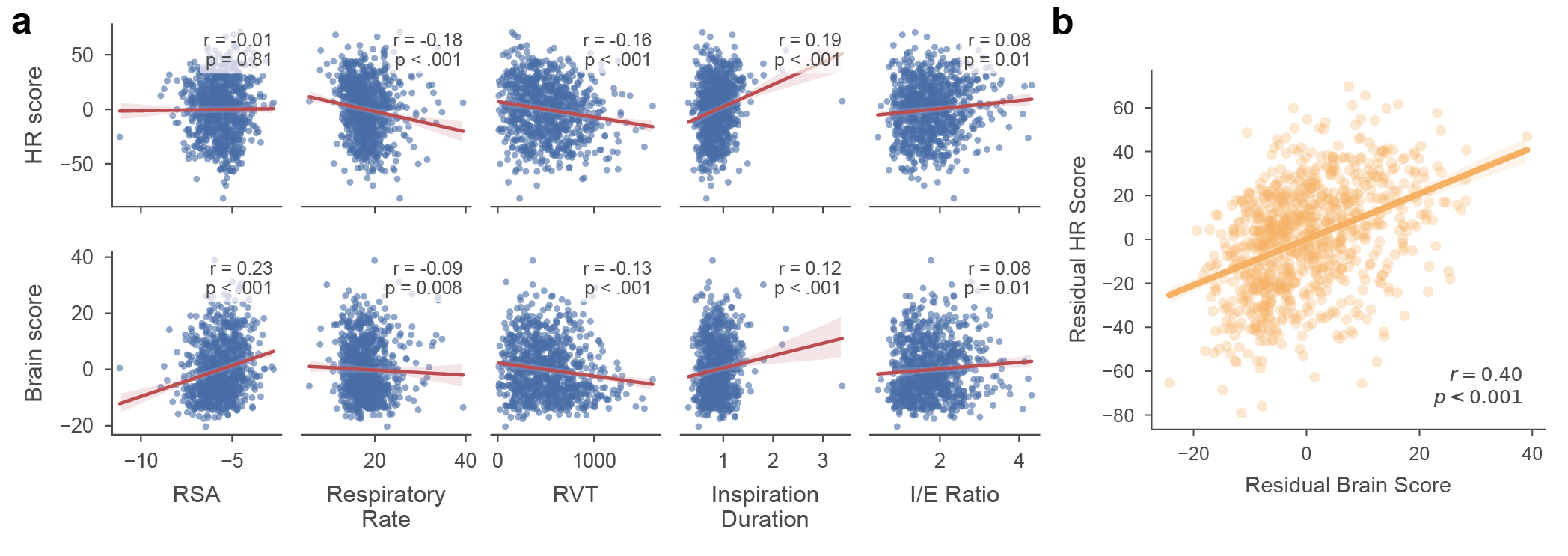

**Figure S3 | Robustness of resting-state heart-brain coupling against respiratory-driven physiological confounds. a**, The pairwise correlations between the PLS HR score (or RSFC score) and each of the five individual respiration-related metrics across subjects. **b**, The correlation between the RSFC scores and HR scores after regressing out respiration-related metrics. The shaded area indicates a 95% confidence interval. *r*, Pearson's correlation coefficient; *p*, *p*-value.

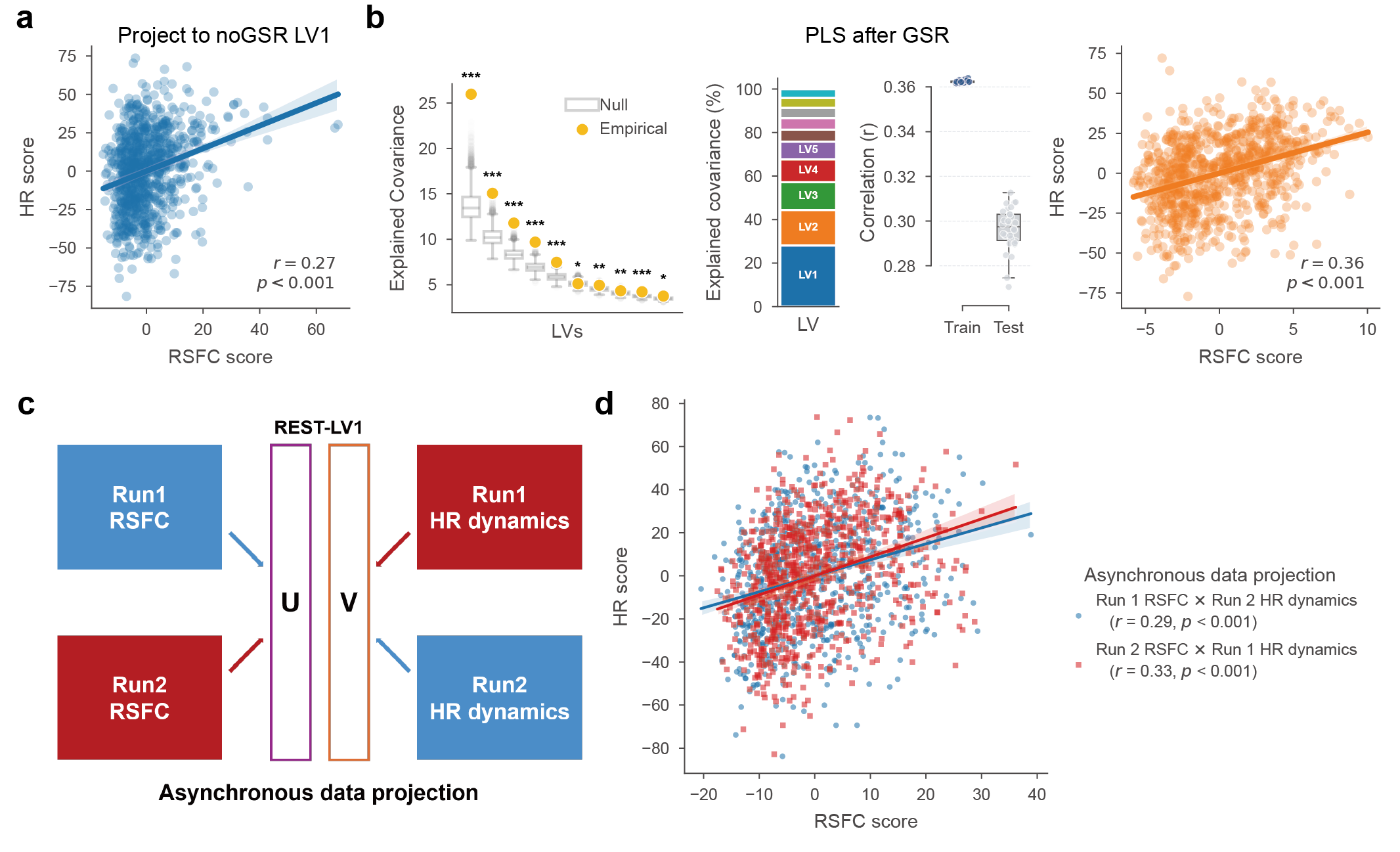

**Figure S4 | Validations of the resting-state heart-brain coupling. a, b,** Evaluation of the PLS model using alternative preprocessing pipelines without global signal regression (noGSR; **a**) and with global signal regression (GSR; **b**). For the GSR pipeline (b), the subpanels present the explained covariance of LVs against 10,000 permutation nulls (left; **p* < 0.05, ***p* < 0.01, ****p* < 0.001), the relative proportion of explained covariance (middle-left), the cross-validation performance (middle-right), and the individual score correlation for LV1 (right). **c, d,** Methodological framework **(c)** and evaluation **(d)** of the asynchronous data projection across separate imaging runs. Panel d presents the generalizability performance, showing the projections of Run 1 RSFC paired with Run 2 HR dynamics (blue) and Run 2 RSFC paired with Run 1 HR dynamics (red).

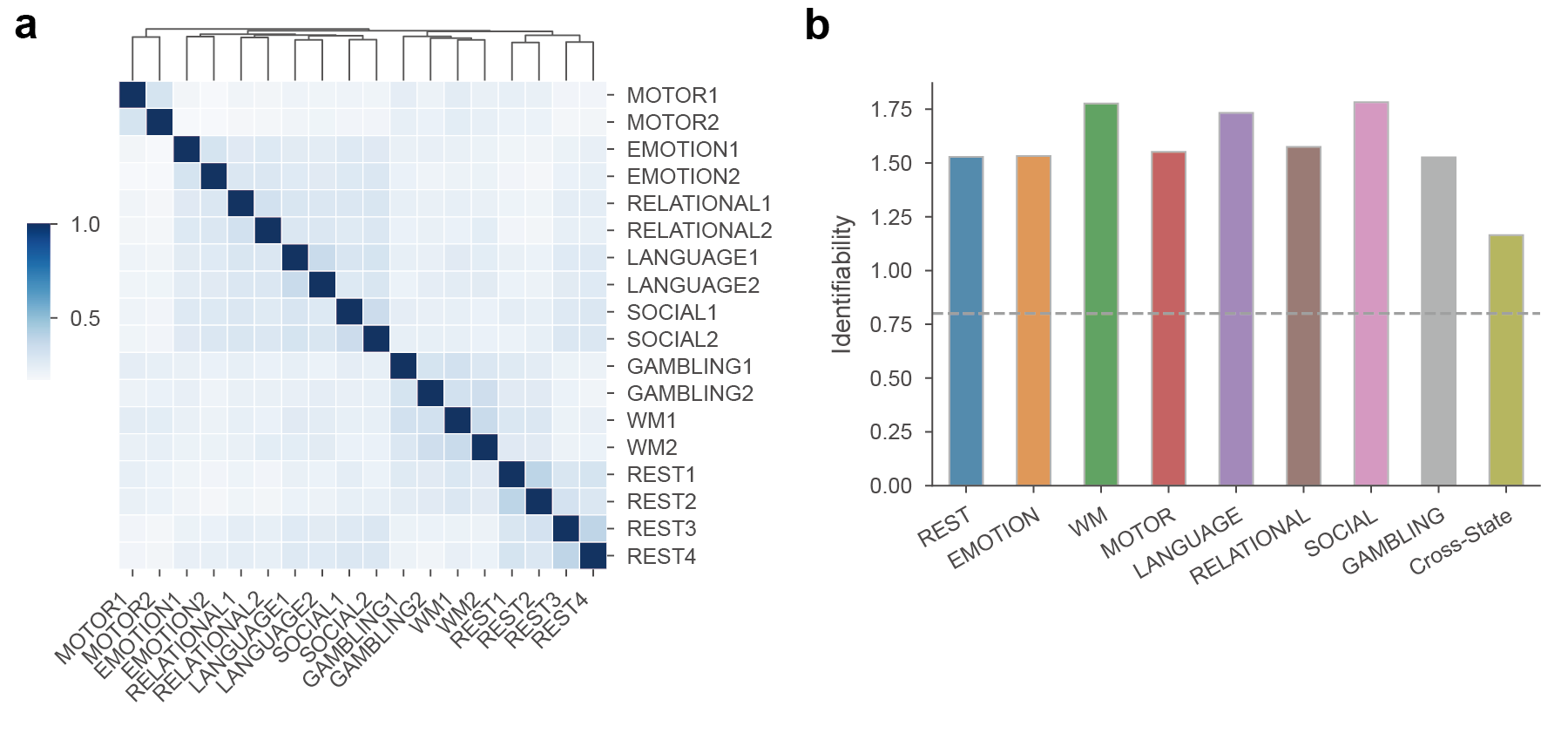

**Figure S5 | Individual specificity and task-state specificity of high-dimensional heart rate dynamics. a**, Pearson correlation matrix displaying the similarity of heart rate dynamics across runs within the same task state or across different task states. **b**, Identifiability of heart rate dynamics evaluated separately for resting-state, task-states, and cross-state.

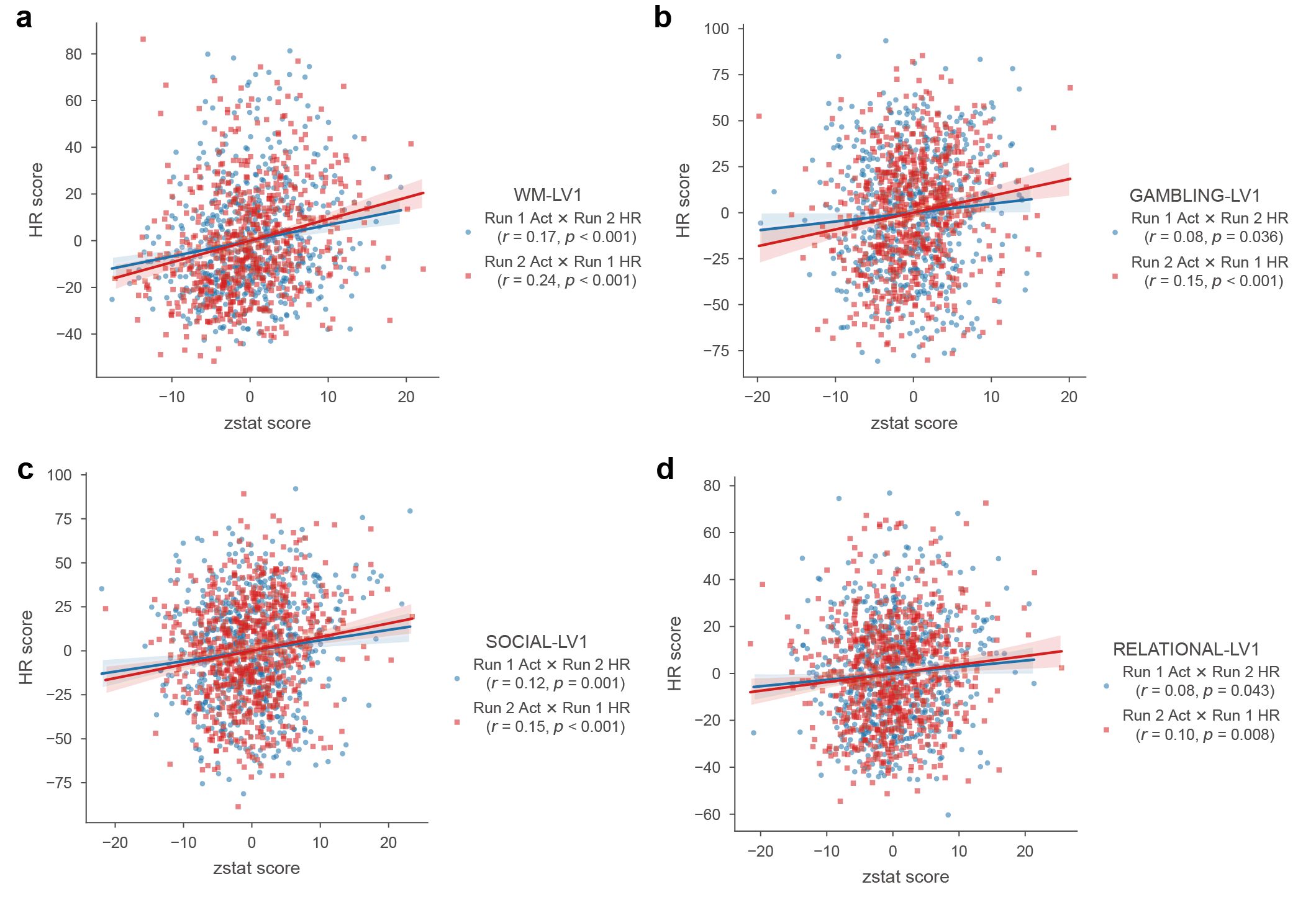

**Figure S6 | Asynchronous cross-run stability of heart-brain covariance during active cognitive states.** The correlation between projected brain and physiological scores for task PLS models under cross-run validation. *r*, Pearson's correlation coefficient; *p*, *p*-value.

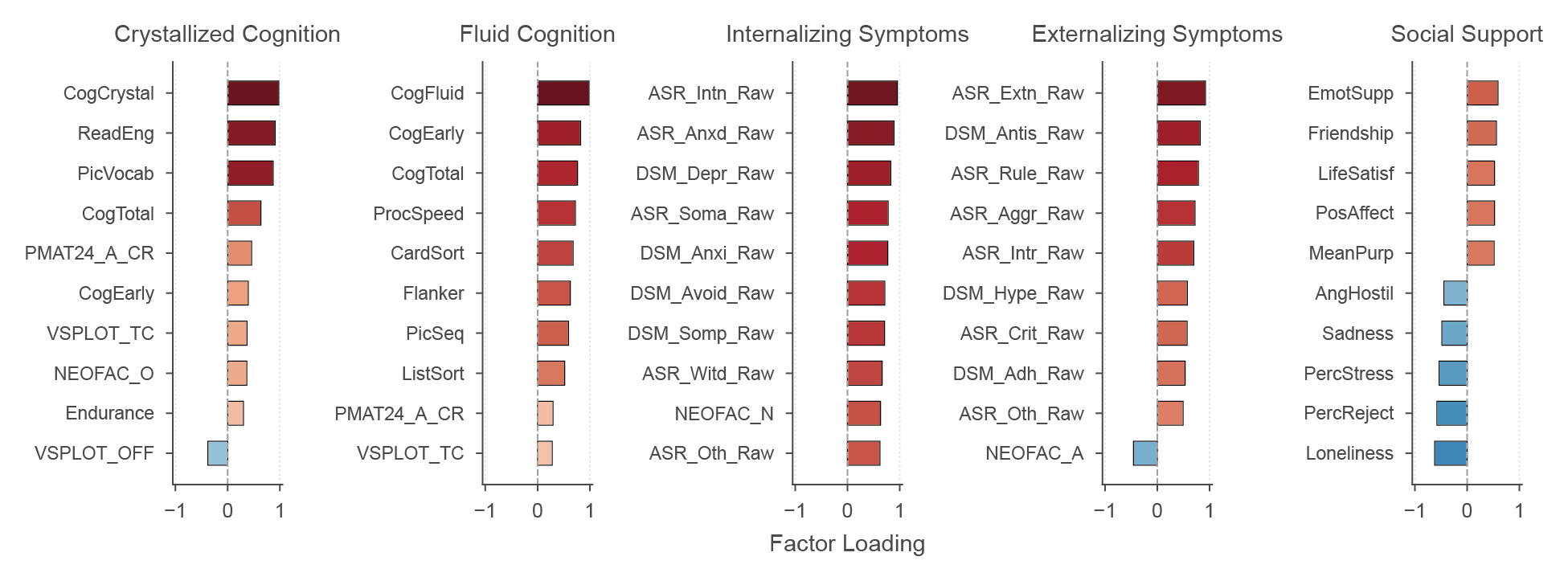

**Figure S7 | Behavioral factors loadings.** The factor loadings of individual behavioral assessments on the five distinct latent factors derived from factor analysis: Crystallized Cognition, Fluid Cognition, Internalizing Symptoms, Externalizing Symptoms, and Social Support.

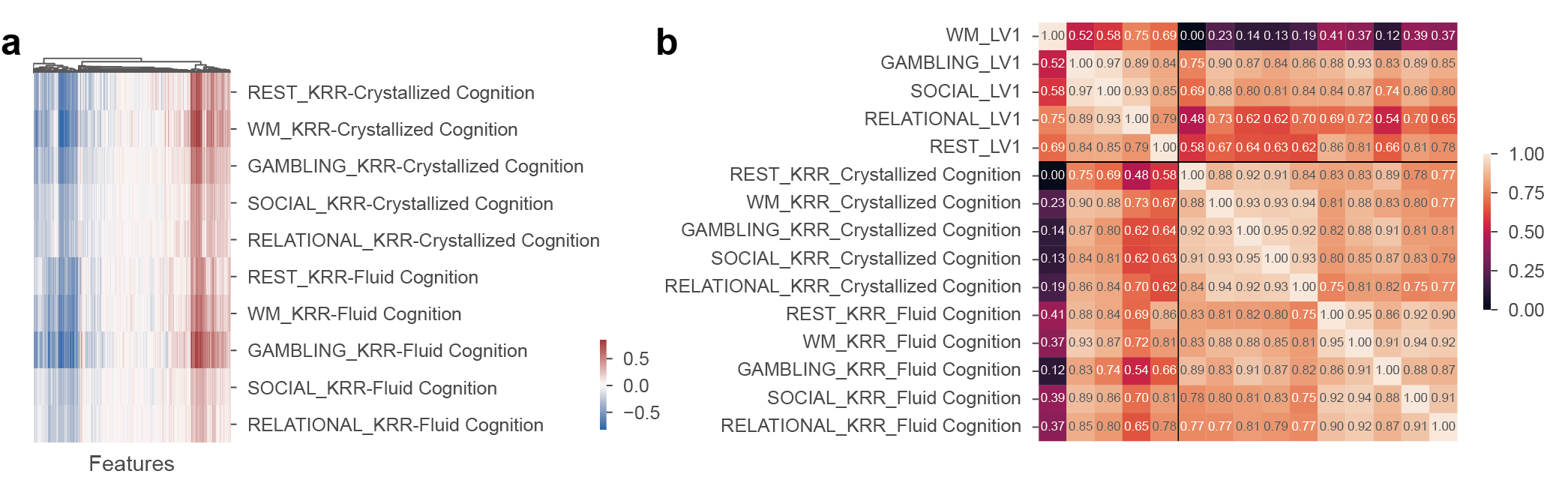

**Figure S8 | Comparison of cognitive predictive heart rate features and latent variable loadings. a**, Haufe activation patterns of heart rate features across different cognitive predictions and brain states. **b**, The similarity of the reconstructed Haufe activation patterns for cognitive predictions and latent variable loadings across resting and task states. The correlation matrix displays Pearson correlation coefficients among the loading or activation pattern vectors.
